# Apoptotic Quality Control of Cellular Competence Limits Cryptic Germinal Center B Cell States

**DOI:** 10.64898/2026.09.10.750694

**Authors:** Chen-Hao Yeh, Nathan Shenkerman, Shengli Song, Hélène Kirshner, Akiko Watanabe, Dongmei Liao, Xiaoe Liang, Wenli Zhang, Kevin Wiehe, Garnett Kelsoe, Masayuki Kuraoka

## Abstract

Germinal center (GC) B cells undergo extensive apoptosis, but how this process shapes evolving GC populations remains unclear. Using mice lacking intrinsic apoptotic effectors BAK and BAX in B cells, we found that mitochondrial apoptosis was dispensable for affinity maturation but critical for controlling GC size and cellular composition, showing that GC population dynamics and affinity maturation can be uncoupled. BAK/BAX-deficient mice mounted enlarged and prolonged GC responses, largely due to the accumulation of non-proliferating Ki-67^lo^CD71^lo^ dark zone (DZ) B cells. CD71^hi^ and CD71^lo^ DZ cells exhibited similar B-cell receptor (BCR) avidity, retained potential for proliferation and plasmacytic differentiation, and shared clonal lineages, indicating recurrent emergence of non-proliferating DZ populations. A subset also carried Somatic-hypermutation-derived defects that abolished antibody expression despite apparently productive V(D)J sequences; reversion of a single amino acid substitution often restored expression. Mitochondrial apoptosis thus enforces cellular quality control in the GC by limiting the persistence of non-proliferating and expression-defective states while remaining dispensable for affinity maturation.

## Introduction

Germinal centers (GCs) are specialized microenvironments in which antigen-specific B cells undergo somatic evolution to generate high-affinity antibodies ^1–8^. During GC reactions, activated B cells undergo iterative rounds of mutations, selection, and proliferation through cycling between light zone (LZ) and dark zone (DZ) that coordinate selection and diversification ^9–14^. In the LZ, B cells acquire cognate antigen displayed on follicular dendritic cells ^15^ through the B cell antigen receptor (BCR) and present peptide-MHCII complexes to T follicular helper (Tfh) cells ^16^. Positively selected B cells subsequently re-enter the DZ for proliferation and diversification before returning to the LZ for additional rounds of selection ^17^, while B cells fail to receive sufficient selection signals undergo apoptosis and are subsequently cleared by tingible body macrophages ^18^.

BCR avidity (or affinity) for cognate antigen is a key driver of GC somatic evolution. B cells expressing higher-affinity BCRs receive enhanced survival signals through BCR engagement with cognate antigen ^19,20^. In addition, higher-affinity B cells capture more antigen and present greater amounts of peptide-MHCII complexes to Tfh cells. B cells that received enhanced T cell help preferentially expand within the DZ before re-entry into the LZ for subsequent rounds of selection ^21–24^. However, GC selection permits persistence of B cells spanning a broad range of BCR affinity and preserving clonal diversity ^25–27^, suggesting that GC evolution is not driven solely by stringent affinity threshold.

B cells undergo extensive proliferation and cell death in GCs. Resulting rapid population turnover accelerates somatic evolution. Proliferating DZ B cells undergo somatic hypermutation (SHM) at immunoglobulin (Ig) loci through the activity of activation-induced cytidine deaminase (AID) ^28^ and generate clonal descendants that express mutated B cell receptors (BCRs) ^3,29–31^. Although SHM preferentially targets hotspot motifs within Ig variable regions^32^, its effects on BCR affinity and receptor integrity are inherently stochastic. Most mutations reduce affinity rather than improve it, and some mutations directly impair BCR expression or function by introducing premature stop codons, frameshift insertions or deletions, or disrupting BCR structure or Ig chain pairing ^33–35^. Therefore, GC reactions continuously generate large numbers of B cells that carry nonproductive or functionally debilitating BCRs. Although apoptotic cell death has been implicated in shaping clonal selection during affinity maturation ^5,20,36–38^, these observations suggest that it may play broader roles in maintaining the integrity and composition of evolving GC populations.

To assess roles for mitochondrial apoptosis during GC responses, we used mice in which B cells lack both BAK and BAX (BAKBAX^B-null^ mice) ^39,40^. BAK and BAX are pro-apoptotic members of the BCL-2 family that mediate mitochondrial outer membrane permeabilization during intrinsic apoptosis ^41,42^. In response to apoptotic stimuli, activated BAK and BAX oligomerize to form pores on the mitochondrial outer membrane. The formation of these pores triggers release of cytochrome c and SMAC and subsequent activation of the caspase cascade. In the absence of both BAK and BAX, B cells become highly resistant to diverse apoptotic stimuli ^39,43^.

Here, we show that immunization of BAKBAX^B-null^ mice elicited markedly enlarged and prolonged GC responses. These enlarged GCs supported B cell affinity maturation comparable to that in B6 GCs, demonstrating that GC population dynamics can be uncoupled from affinity maturation. Enlarged GCs in BAKBAX^B-null^ mice accumulated a distinct population of Ki-67^lo^CD71^lo^ DZ B cells that retained proliferative capacity and potential for plasmacytic differentiation. CD71^lo^ and CD71^hi^ DZ B cells exhibited comparable BCR avidity for cognate antigen and accumulated similar levels of V(D)J mutations. Coexistence of CD71^lo^ and CD71^hi^ DZ B cells within individual clonal lineages indicated that non-proliferating DZ cells recurrently emerge during GC responses. V_H_ gene segments that were enriched in CD71^lo^ DZ populations were subsequently depleted during the GC responses, indicating the link between the non-proliferating DZ state and declining clonal fitness. A substantial fraction of BAK/BAX-deficient DZ cells expressed BCRs carrying non-canonical debilitating mutations that preserved apparent sequence integrity but impaired antibody expression. These findings show that mitochondrial apoptosis constrains the size and composition of evolving GC populations by limiting the accumulation of cryptic non-proliferating populations and cells carrying SHM-derived defects in antibody expression. This function is not required to maintain affinity maturation. Together, our observations identify cellular competence as an underappreciated dimension of clonal fitness during GC evolution.

## Results

### Apoptosis in B cells controls magnitude and duration of GC responses

To assess the contribution of B cell apoptosis to GC responses, we immunized B6, *Bak*^-/-^Bax^fl/fl^ (BAK^null^), and *Cd19*^Cre/wt^*Bak*^-/-^*Bax*^fl/fl^ (BAKBAX^B-null^) mice in a hind footpad with recombinant protective antigen (rPA) of *Bacillus anthracis* in alhydrogel. We monitored GC responses in the draining popliteal lymph node (LN) for up to 40 days (**Fig. 1**). In B6 mice, both the frequency and number of B220^+^CD138^-^GL7^+^CD38^lo^IgD^-^ GC B cells peaked between day 12 and day 16, waned to ∼30% of the peak response by day 24, and remained relatively stable through day 40 (**Fig. 1**). Both the kinetics and magnitude of GC responses were comparable between B6 and BAK^null^ mice (*p* > 0.45), indicating that loss of BAK alone has minimal impact on GC responses.

**Figure 1.**
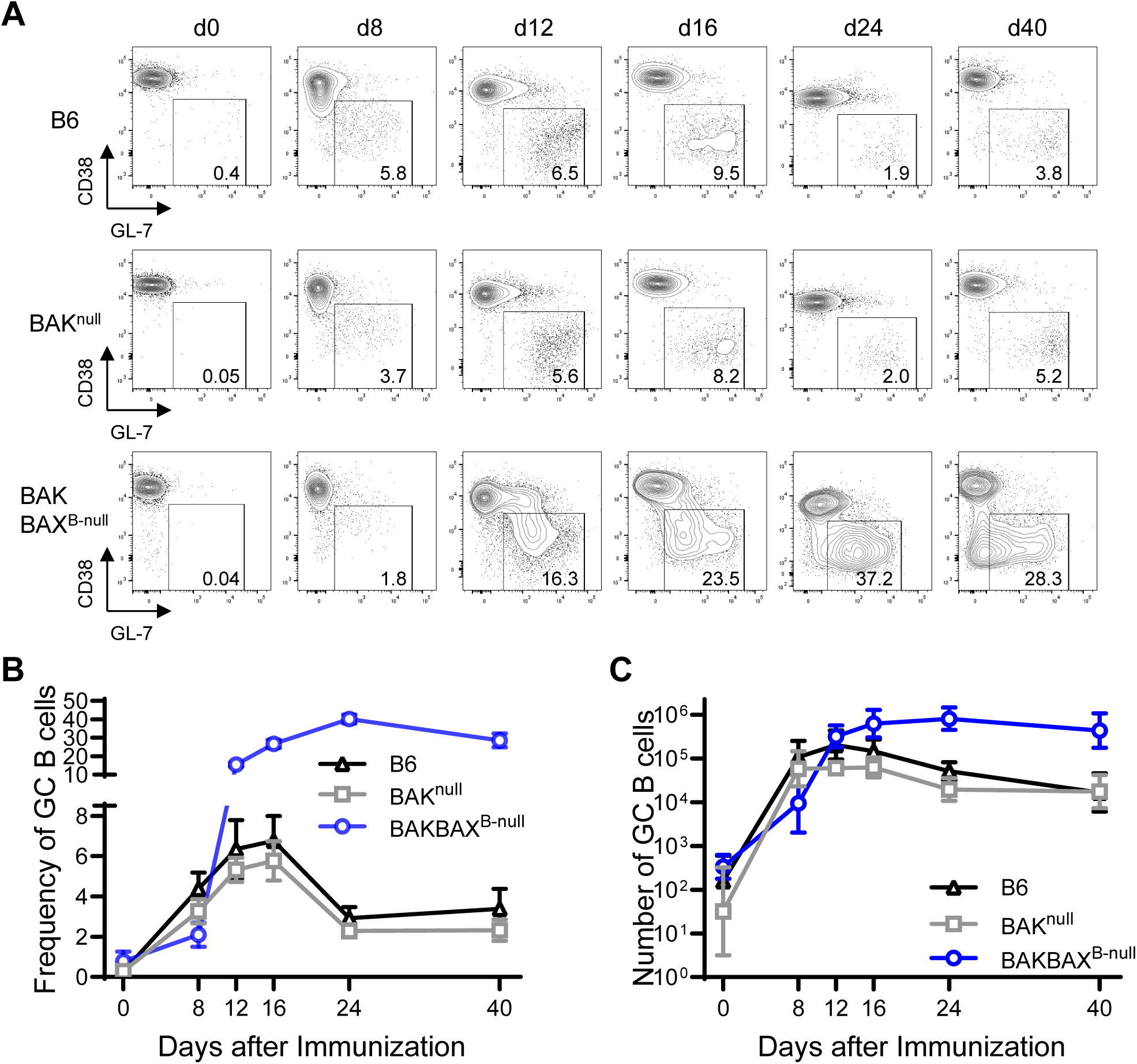
BAKBAX^B-null^ mice mount robust GC responses following rPA immunization. (**A**) Representative flow diagrams of GL-7 and CD38 expressions by B220^+^CD138^-^ cells in the draining LNs of B6, BAK^null^, and BAKBAX^B-null^ mice following footpad immunization with rPA in alhydrogel. Numbers within boxes represent frequency of GL-7^+^CD38^lo^ cells among B220^hi^CD138^-/lo^ cells. (**B** and **C**) Kinetics of frequency and number of GC B cells in the draining LNs of B6 (n = 4 – 12), BAK^null^ (n = 4 – 8), and BAKBAK^B-null^ mice (n = 5 – 13). (**B**) Frequency of GL-7^+^CD38^lo^IgD^-^ GC B cells among B220^+^CD138^-^ cells. Mean values for each time point are connected. Error bars, SD. (**C**) Geometric mean values for each time point are connected. Error bars, geometric SD. Combined data from at least 2 independent experiments for each time point are shown. Cell counts were log10-transformed prior to statistical analysis. Statistical significance was tested using a two-way Mixed-effects model.

In contrast, GC responses differed significantly in BAKBAX^B-null^ mice (*p* < 0.001 versus B6 or BAK^null^ mice; **Fig. 1**). Day 8 GC responses in BAKBAX^B-null^ mice were smaller than those in B6 controls (∼2-fold and ∼5-fold lower in frequency and cell number, respectively). This initially smaller GC response is consistent with impaired proliferative responses of BAK/BAX-deficient B cells to mitogenic stimuli ^39^. By day 12, GC B cell frequencies in BAKBAX^B-null^ mice were ∼2-fold higher than those in B6 mice (*p* < 0.01). GC B cell numbers were not significantly different between B6 and BAKBAX^B-null^ mice at this time point. By day 16, both the frequency and number of GC B cells were ∼4-fold higher in BAKBAX^B-null^ mice (*p* < 0.001). In contrast to B6 GC responses, which substantially contracted after day 16, GC responses in BAKBAX^B-null^ mice continued to expand through day 24 and remained elevated. BAK/BAX-dependent apoptosis constrains both the magnitude and persistence of GC responses.

### GC B cells in B6 and BAKBAX^B-null^ mice exhibit similar V_H_ repertoires and SHM

To determine whether enlarged GC responses in BAKBAX^B-null^ mice altered dynamics of GC B cell repertoires, we performed single-cell Nojima cultures on GC B cells sorted at days 8, 16, and 24 following rPA immunization ^25,44^. We amplified IgH V(D)J rearrangements from a selected subset of cultures that contained clonal IgGs. Early GC responses in B6 and BAKBAX^B-null^ mice showed similar V_H_ gene usage. From day 8 GCs, we recovered V_H_14-4, V_H_1-82, and V_H_8-8 gene segments at high frequency, accounting for approximately half of recovered V_H_ gene segments (**Fig. 2A** and **Table S1**) ^25^. As GC responses progressed, these initially dominant V_H_ gene segments contracted substantially and were replaced by previously rarer V_H_ clones in both B6 and BAKBAX^B-null^ mice. Overall, B cell-specific deletion of BAK/BAX-dependent apoptosis did not substantially alter recruitment or diversification of V_H_ repertoires during GC responses.

**Figure 2.**
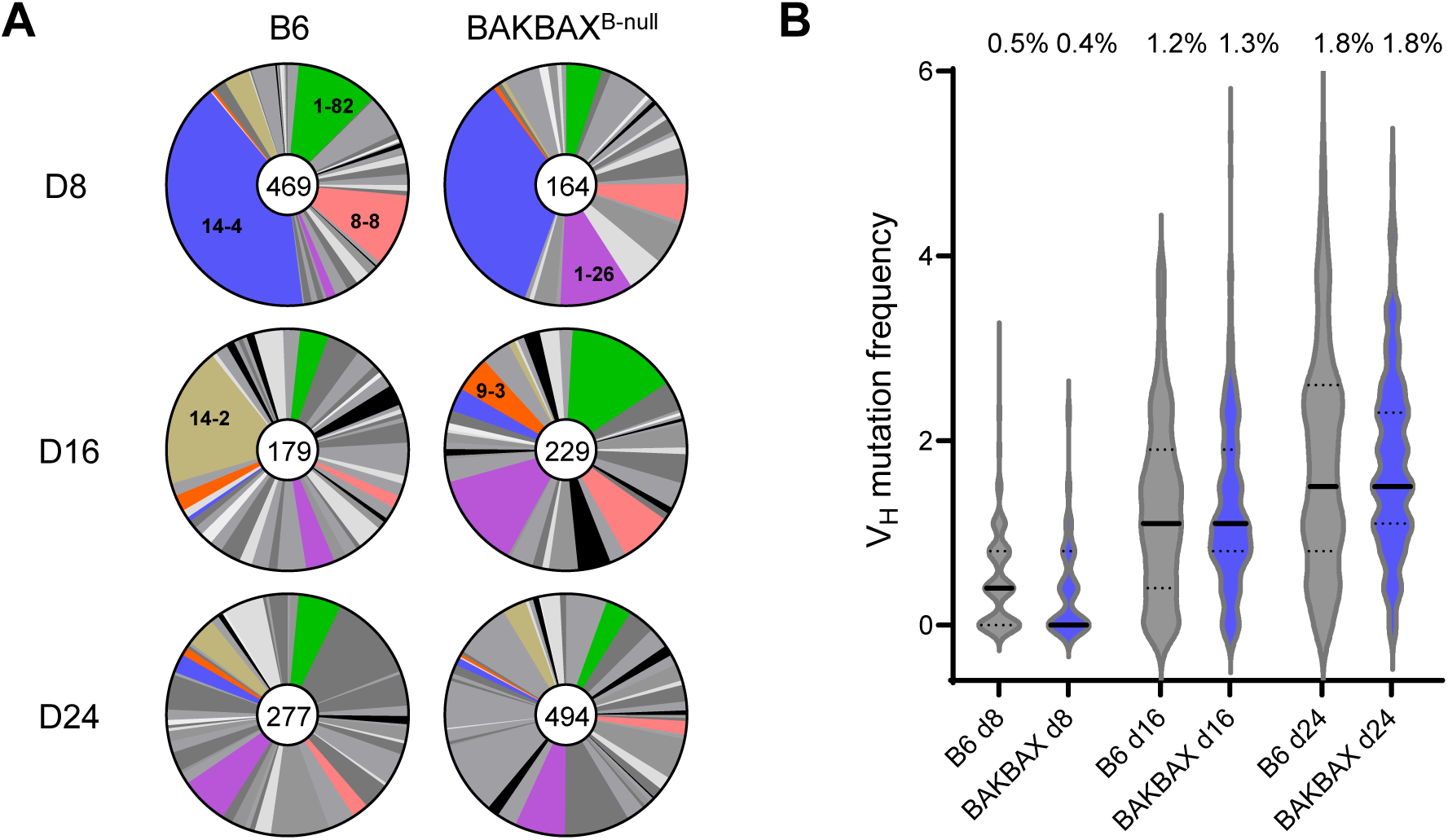
Somatic genetics of GC B cells in B6 and BAKBAX^B-null^ mice. Rearranged VDJ sequences were obtained from clonal cultures of day 8, 16, and 24 GC B cells from B6 and BAKBAX^B-null^ mice. (**A**) Pie charts depict proportion of V_H_ gene segments. Numbers in middle circles represent number of V_H_ gene segments analyzed. (**B**) Violin plots show distribution of V_H_ mutation frequency of GC B cells in (**A**). Horizontal solid lines, median. Dotted lines, quartiles. Percentage values at the top of the figure indicate average mutation frequency for each group. Mutation frequencies represent number of nucleotide substitutions per base pairs sequenced for each V_H_ gene. Statistical significance was tested by Kruskal-Walis test with Dunn’s multiple comparisons. No statistical significance between B6 and BAKBAX GCs for all time points. Mutation frequencies significantly increased from d8 to d24 GCs for both B6 and BAKBAX^B-null^ mice (*p* < 0.001).

GC B cells from B6 and BAKBAX^B-null^ mice accumulated V_H_ point mutations at similar rates throughout the GC response (**Fig. 2B**). Average V_H_ mutation frequencies of GC B cells increased from day 8 (∼0.5 × 10^-2^) to day 16 (∼1.2 × 10^-2^) and day 24 (∼1.8 × 10^-2^). The proportion of mutated GC B cells also increased from day 8 (∼55%) to day 24 (∼94%). Nearly half of day 24 GC B cells carried ≥5 V_H_ point mutations, whereas only 2% -3% of day 8 GC B cells did. Ablation of BAK/BAX-dependent apoptosis neither enhanced nor impaired accumulation of V_H_ SHM during GC responses.

### Enlarged GCs in BAKBAX^B-null^ mice support affinity maturation

Enlarged GC responses in BAKBAX^B-null^ mice could reflect persistence of unspecific or low-affinity B cells that would normally be eliminated during GC competition. If so, antigen-specific GC B cells would be less frequent and exhibit lower BCR avidities in BAKBAX^B-null^ mice. To assess BCR specificity and avidity of GC B cells, we used single-cell Nojima cultures. We generated total 7,689 IgG^+^ clonal cultures from B6 (n = 3,409) and BAKBAX^B-null^ GC B cells (n = 4,280) following immunization with rPA as in **Fig. 1** and **Fig. 2**. We determined specificity and avidity of culture supernatant IgGs by a multiplex Luminex assay. Among B6 GC B cell cultures, frequencies of rPA-binding clonal IgGs among all clonal IgGs increased from day 8 (∼14%) to day 12 (∼33%) and remained elevated at day 16 (∼38%) (**Fig. 3A**). These frequencies dropped at day 24 (∼27%) and remained at similar levels at day 40 (∼25%). In BAKBAX^B-null^ mice, frequencies of rPA-binding clonal IgGs were similar to or higher than those in B6 mice throughout the GC responses (**Fig. 3A**). Frequencies of rPA-binding clonal IgGs increased from day 8 (∼16%) to day 12 (∼46%) and remained elevated at all later timepoints (∼47-53%) (**Fig. 3A**).

**Figure 3.**
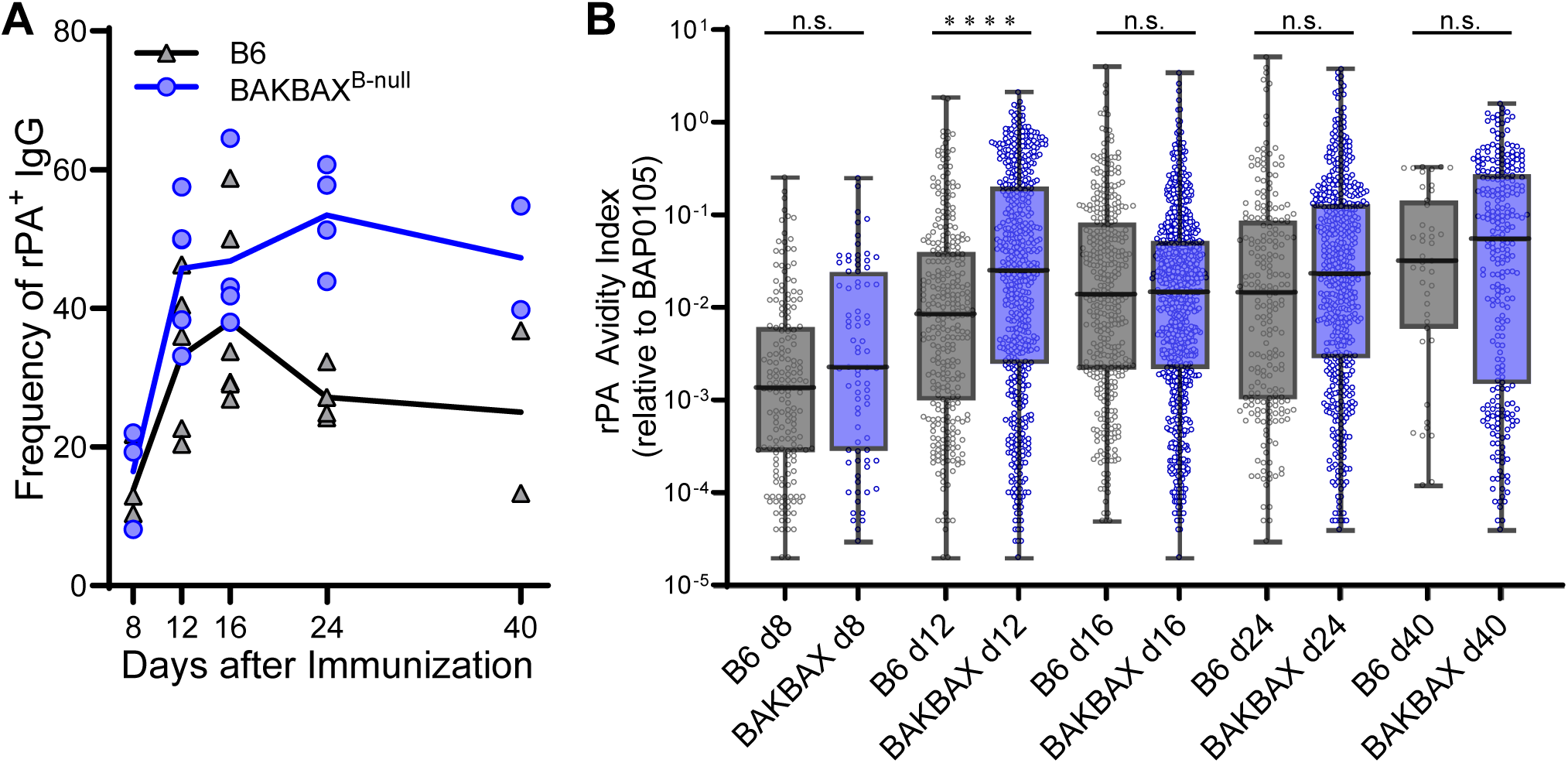
Enlarged GC responses in BAKBAX^B-null^ mice support normal affinity maturation. (**A**) Frequencies of rPA-binding clonal IgGs among all clonal IgGs are shown. Each symbol represents each mouse (n = 2 – 6). Mean values of each timepoint are connected with lines for each mouse strain. (**B**) Distribution of AvIn values of clonal IgGs from individual, single B cell cultures. Each dot represents an AvIn value for an individual clonal IgG sample (n = 43 – 848). Boxes extend from the 25th to 75th percentiles and lines in the boxes represent medians. Error bars represent minimum to maximum values. ****, *p* < 0.0001; n.s., *p* > 0.05 by Kruskal-Walis test with Dunn’s multiple comparisons.

To determine the distribution of BCR avidities for rPA, we calculated avidity index (AvIn) value for each clonal IgG in reference to a high-affinity rPA-specific monoclonal antibody BAP0105 ^25^. Among B6 GC B cell cultures, geometric mean AvIn values for rPA-binding clonal IgGs increased ∼10-fold from day 8 (AvIn = 0.001) to day 16 (AvIn = 0.01) and plateaued thereafter (**Fig. 3B**). The rPA-binding clonal IgGs from BAK/BAX-deficient GC B cells exhibited similar levels of AvIn values at all matched time points, except for day 12. On day 12, geometric mean AvIn value was higher in BAK/BAX-deficient GC B cells than in B6 GC B cells (AvIn = 0.026 vs. 0.009). Enlarged and prolonged GC responses in BAKBAX^B-null^ mice retained antigen-specific B cells and supported affinity maturation as efficiently as those in B6 mice.

### BAKBAX^B-null^ GCs accumulate non-proliferating DZ populations

To further characterize B cell populations in enlarged GCs in BAKBAX^B-null^ mice, we performed high-dimensional flow cytometric analysis and identified marked expansion of phenotypically distinct DZ populations (**Fig. 4**). In B6 mice, the ratio of CXCR4^+^CD86^lo^ DZ cells to CXCR4^-^CD86^hi^ LZ cells was approximately 2.8:1. In contrast, the DZ to LZ ratio in BAKBAX^B-null^ mice was approximately 10:1, indicating substantial enrichment of DZ cells (**Figs. 4A** and **4B**). Uniform manifold approximation and projection (UMAP) analysis identified conventional LZ, DZ1 (BCR^hi^Ki-67^hi^), and DZ2 (BCR^lo^Ki-67^hi^) populations in both B6 and BAKBAX^B-null^ mice, whereas distinct populations of DZ3 (BCR^hi^Ki-67^lo^) and DZ4 (BCR^lo^Ki-67^lo^) were selectively present in BAKBAX^B-null^ mice (**Fig. 4C**). Consistent with UMAP analysis, more than 95% of DZ cells in B6 mice expressed cell proliferation markers (Ki-67^hi^ or CD71^lo^), whereas non-proliferating DZ cells were readily detectable in BAKBAX^B-null^ mice (**Fig. 4D**). Ki-67 and surface BCR expressions by LZ cells were comparable between B6 and BAKBAX^B-null^ mice. These findings indicate that BAK/BAX-dependent apoptosis normally constrains accumulation of non-proliferating DZ populations during GC responses.

**Figure 4.**
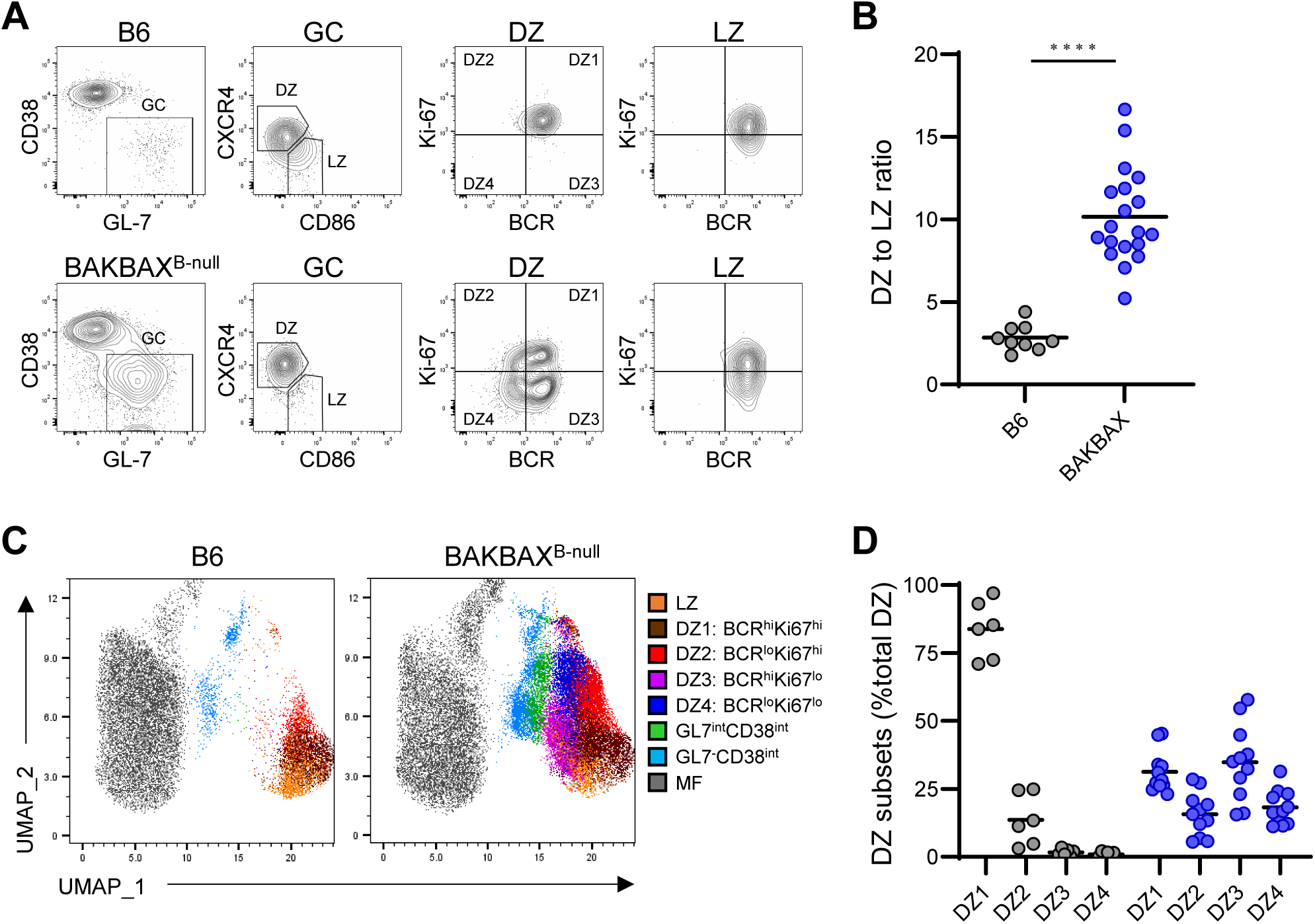
Enlarged GCs in BAKBAX^B-null^ mice accumulate non-proliferating DZ cells. B6 and BAKBAX^B-null^ mice were immunized with rPA in alhydrogel, and GC responses were analyzed at day 16 following immunization. (**A**) Representative flow diagrams of GC, DZ (CD86^lo^CXCR4^hi^) and LZ (CD86^hi^CXCR4^lo^) cells, and surface BCRs (determined by Igκ and Igλ expressions) and intracellular Ki-67 expressions by DZ and LZ cells are shown. (**B**) DZ to LZ ratio for B6 and BAKBAX-deficient GC B cells are shown. Each dot represents an individual mouse. Horizontal bars represent mean. ****, *p* < 0.0001 by unpaired *t* test with Welch’s correction. (**C**) UMAP plots. Specific B cell subsets were colored according to key. (**D**) Distributions of DZ subsets among all DZ cells in B6 (black) or in BAKBAX^B-null^ mice (blue). DZ1 – DZ4 subsets were defined based on combinations of BCR and Ki-67 expression as follows: DZ1, BCR^hi^Ki-67^hi^; DZ2, BCR^lo^Ki-67^hi^; DZ3, BCR^hi^Ki-67^lo^; and DZ4, BCR^lo^Ki-67^lo^. In experiments incompatible with intracellular Ki-67 staining, surface CD71 expression was used as a surrogate marker for Ki-67 to define corresponding DZ subsets. Each dot represents an individual mouse. Horizontal bars represent mean. Combined data from 4 independent experiments are shown.

### Non-proliferating DZ cells retain antigen-binding BCR avidity and proliferative potential

Non-proliferating DZ cells in BAKBAX^B-null^ mice might represent B cells that failed affinity-dependent competition during GC responses. If so, BCR avidities of non-proliferating DZ cells would be lower than those of proliferating DZ cells. To compare BCR specificity and avidity between proliferating and non-proliferating DZ populations, we used single-cell Nojima cultures. Because intracellular Ki-67 staining requires fixation and permeabilization steps that are incompatible with Nojima cultures, we used CD71 as a surrogate surface marker for intracellular Ki-67 ^45^. Consistent with a previous report ^45^, CD71 expression strongly correlated with intracellular Ki-67 expression among GC B cells in both B6 and BAKBAX^B-null^ mice (**Fig. S1**). We therefore sorted CD71^hi^ and CD71^lo^ DZ cells from day 16 PLNs of rPA-immunized BAKBAX^B-null^ mice for Nojima culture analysis (CD71^hi^BCR^all^ and CD71^lo^BCR^all^ cells; **Fig. 5A**). In some experiments, we subdivided CD71^hi^ and CD71^lo^ DZ cells into BCR^hi^ and BCR^lo^ populations prior to culture (**Fig. 5B**). In total, we generated 1,989 IgG^+^ clonal cultures for analysis.

**Figure 5.**
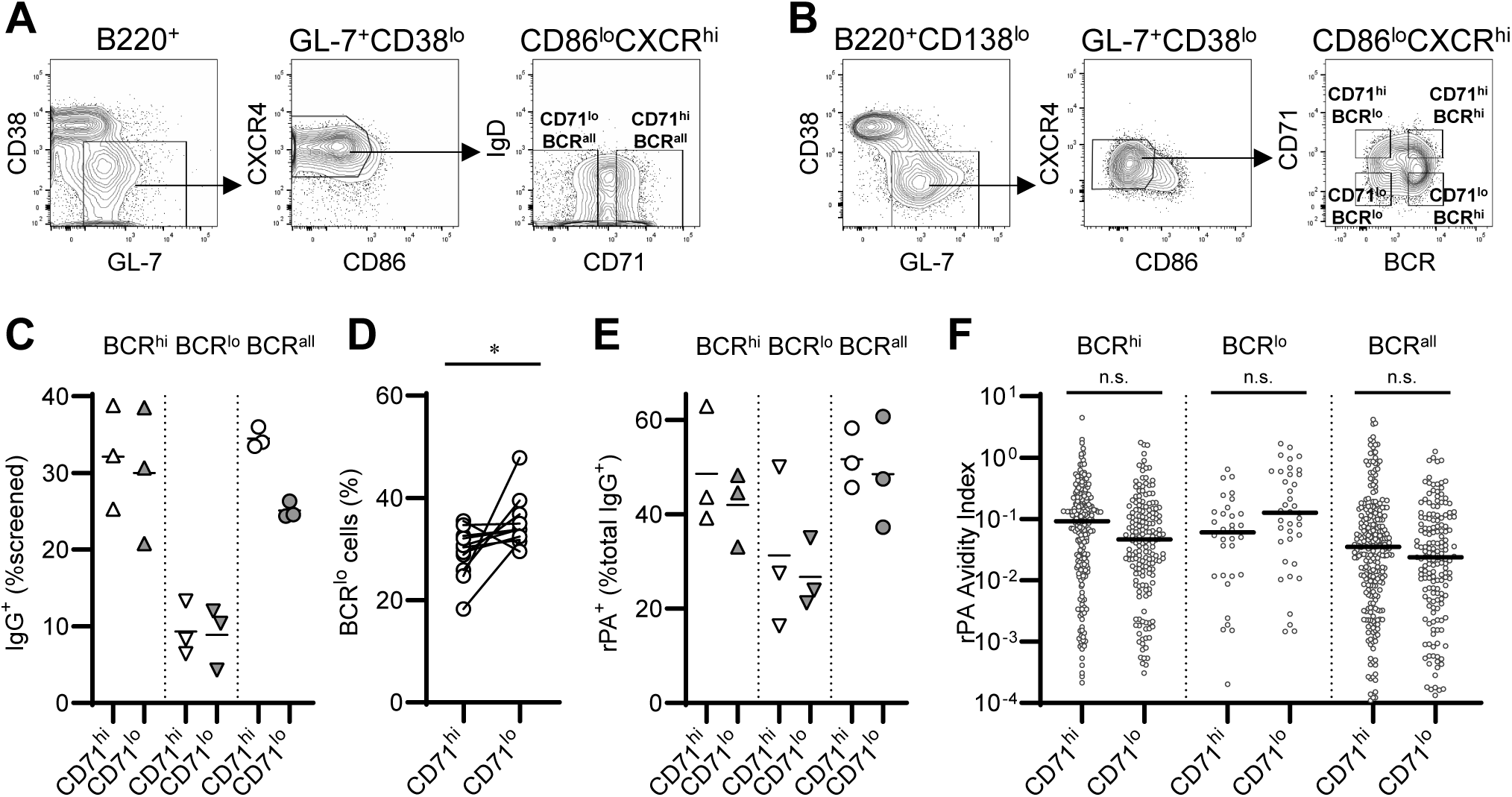
BCR specificity and avidity of non-proliferating DZ cells. CD71^hi^ or CD71^lo^ DZ cells were sorted from draining LNs of rPA-immunized BAKBAX^B-null^ mice (day 16) for Nojima cultures. (**A**) CD71^hi^ and CD71^lo^ DZ cells were sorted not considering surface BCR expression (BCR^all^). (**B**) CD71^hi^ and CD71^lo^ DZ cells were further separated by surface BCR expression levels. (**C**) Cloning efficiency (Number of IgG positive samples / Number of GC B cell samples screened). Each symbol represents an individual mouse. (**D**) Frequencies of BCR^lo^ cells in CD71^hi^ or CD71^lo^ cells are shown. *, *p* < 0.05 by Wilcoxon matched-pairs signed rank test. (**E**) Frequency of rPA-specific IgGs among all clonal IgGs. Each symbol represents an individual mouse. (**F**) Distribution of AvIn values of culture supernatant IgGs from individual, single B cell cultures. Each dot represents an AvIn value for an individual clonal IgG sample (n = 38 – 252). n.s., *p* > 0.05 by Kruskal-Walis test with Dunn’s multiple comparisons. Combined data from 3 independent experiments are shown.

Cloning efficiency, defined as the proportion of IgG^+^ cultures, was comparable between CD71^hi^ and CD71^lo^ DZ B cells when sorted according to BCR expression (**Fig. 5C**). Recovery of IgG^+^ cultures from CD71^lo^ DZ cells indicates that CD71^lo^ DZ cells retain capacities for proliferation and differentiation into antibody-secreting cells despite their non-proliferating state *in vivo*. Notably, cloning efficiencies of BCR^lo^ cells were lower than those of BCR^hi^ cells regardless of CD71 expression. Cloning efficiency was moderately lower for CD71^lo^BCR^all^ cells that contained BCR^lo^ cells at elevated frequency (**Figs. 5C** and **5D**).

We compared BCR specificity and avidity between CD71^hi^ and CD71^lo^ DZ cells. Frequencies of rPA-binding clonal IgGs were comparable between CD71^hi^ and CD71^lo^ DZ cells within BCR^hi^, BCR^lo^, and BCR^all^ populations (**Fig. 5E**). Distributions of rPA AvIn values did not differ between CD71^hi^ and CD71^lo^ DZ cells (**Fig. 5F**). Non-proliferating state was not associated with reduced BCR specificity or avidity; therefore, loss of affinity-dependent competition is unlikely to be the primary driver of entry into the non-proliferating DZ compartment.

### BCR repertoires enriched in CD71^lo^ DZ cells contract during GC progression

CD71^lo^ DZ cells might represent B cell populations that are contracting during GC responses. To investigate the relationship between the CD71^lo^ DZ state and clonal dynamics during GC responses, we obtained paired Ig(H+L) V(D)J sequences from 476 CD71^hi^ and 362 CD71^lo^ DZ cells from BAKBAX^B-null^ mice 16 days following rPA immunization (**Fig. 6A**). The CD71^hi^/CD71^lo^ ratio positively correlated with subsequent changes in V_H_ gene frequency among GC B cells (**Fig. 6B**). Several V_H_ genes enriched in the CD71^lo^ DZ compartment (*e.g.*, V_H_1-82, V_H_8-8, and V_H_9-3) contracted substantially from day 16 to day 24 GC responses (**Figs. 6A** and **6B**). By contrast, V_H_ genes enriched in the CD71^hi^ DZ compartment expanded (*e.g.*, V_H_1-75, V_H_1-85, and V_H_14-2). Notably, the rapid contraction of V_H_14-4 between day 8 and day 16 GC responses (**Fig. 2A**) ^25^ was accompanied by relative enrichment in the CD71^lo^ compartment (**Fig. 6A**). These findings suggest that acquisition of the CD71^lo^ DZ state predicts reduced clonal persistence during GC responses. CD71^lo^ DZ cells may represent descendants of previously expanded GC lineages that are progressively losing competitive fitness.

**Figure 6.**
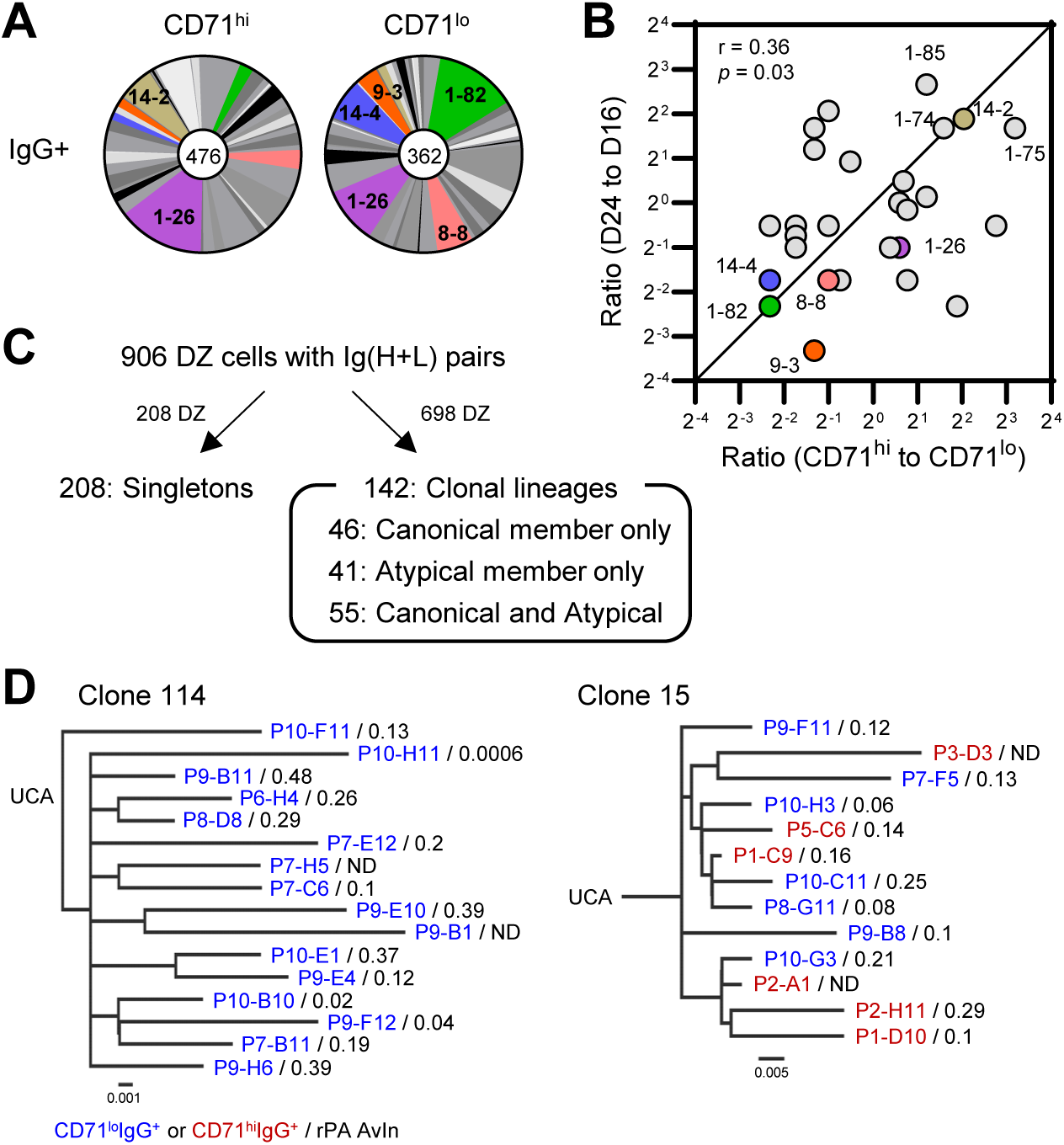
Clonal trajectory of DZ cells in BAKBAX^B-null^ mice. Paired Ig(H+L) V(D)J gene sequences were obtained from CD71^hi^ or CD71^lo^ DZ cells after Nojima culture. (**A**) Pie charts depict proportion of V_H_ gene segments. Numbers in middle circles represent number of V_H_ gene segments analyzed. (**B**) Correlation between preferential representation of V_H_ genes in CD71^hi^ relative to CD71^lo^ DZ compartments and subsequent changes in V_H_ usage during GC progression. The x-axis shows the ratio of V_H_ gene frequencies of CD71^hi^ to CD71^lo^ DZ cells at day 16. The y-axis shows the ratio of V_H_ gene frequencies of day 24 to day 16 total GC B cells (Fig. 2). Each dot represents an individual V_H_ gene. Colored dots indicate representative V_H_ segments discussed in the text. Only V_H_ genes detected in day 16 GC, day 24 GC, CD71^hi^ DZ, and CD71^lo^ DZ populations were included in the analysis. Correlation co-efficient (r) and significance were determined by one-tailed Spearman’s correlation test. (**C**) Clonal lineage analysis of 906 DZ B cells. A total of 208 cells were classified as singletons, whereas the remaining 698 cells formed 142 independent clonal lineages. Among these clonal lineages, 46 consisted exclusively of canonical DZ members (CD71^hi^IgG^+^), 41 consisted exclusively of atypical DZ members (CD71^hi^IgG^-^, CD71^lo^IgG^+^, and/or CD71^lo^IgG^-^), and 55 contained mixtures of both canonical and atypical DZ members. (**D**) Clonal genealogy by V(D)J sequences of CD71^hi^ and CD71^lo^ DZ B cells. Left, Clone 114. Members of this clone use V_H_1-82/D_H_4-1/J_H_4 and V_κ_1-110/J_κ_1 rearrangements. Right, Clone 15. Members of this clone use V_H_1-39/D_H_2-5/J_H_1 and V_κ_19-93/J_κ_2 rearrangements. The rPA AvIn values are indicated for each member of these clones. These phylogenic trees were inferred using Cloanalyst.

### Clonal trajectory of CD71^hi^ and CD71^lo^ DZ cells

Based on similarity in the V_H_ and J_H_ gene usage and in the HCDR3 sequence, we determined clonal relationships among DZ subsets using Cloanalyst ^46^. For this analysis, we also included V(D)J sequences from 68 DZ cells (34 each from CD71^hi^ and CD71^lo^ subsets) that expanded in Nojima cultures without secreting IgG (discussed further below). Among 906 DZ cells for which we obtained Ig(H+L) pair sequences, 208 (23%) were singletons (**Fig. 6C**). The remaining 698 cells (77%) belonged to clonal lineages. We obtained total 142 clonal lineages; 46 (32%) contained CD71^hi^IgG^+^ canonical DZ members only, 41 (29%) contained atypical DZ members only (*i.e*., CD71^hi^IgG^-^, CD71^lo^IgG^+^, and CD71^lo^IgG^-^ DZ cells), and 55 (39%) contained both canonical and atypical DZ members (**Fig. 6C**).

Clonal lineages enriched for non-proliferating DZ members likely represent previously expanded lineages that are undergoing clonal contraction (**Fig. 6B**). For example, clonal lineage 114 consisted entirely of CD71^lo^IgG^+^ DZ cells (**Fig. 6D**). Although 2 of 16 lineage members did not bind rPA, the remaining members exhibited a median AvIn value of 0.2, a value higher than the median AvIn values of bulk DZ populations (**Figs. 5F** and **6D**). Clonal lineages retaining substantial BCR avidity may nevertheless undergo clonal contraction.

Nearly 40% of clonal lineages contained both canonical and atypical DZ members. For example, clonal lineage 15 contained 13 members, including 6 CD71^hi^IgG^+^ and 7 CD71^lo^IgG^+^ DZ cells (**Fig. 6D**). CD71^lo^IgG^+^ members appeared throughout the tree rather than being confined to particular branches (**Fig. 6D**). CD71^hi^ and CD71^lo^ members exhibited median AvIn values at a similar level (0.15 and 0.12, respectively; **Fig. 6D**). Entry into the CD71^lo^ non-proliferating DZ state may frequently occur within clonal lineages without measurable loss of BCR avidity.

### SHM generates non-canonical debilitating immunoglobulin mutations in GC B cells

A subset of DZ cells expanded during Nojima cultures without secreting IgG. We observed IgG^-^ cultures among CD71^hi^ and CD71^lo^ BAKBAX^B-null^ DZ cells but these cultures were rare among B6 GC B cells (**Table S2**). To characterize IgG^-^ DZ cells, we compared V(D)J sequences between IgG^-^ and IgG^+^ DZ cells. While V_H_ and V_K/L_ mutation frequencies were comparable between IgG^-^ and IgG^+^ DZ cells (**Fig. 7A**), V(D)J sequences from IgG^-^ DZ cells exhibited elevated R/S (replacement/silent) mutation ratios within V_H_ framework regions compared with IgG^+^ DZ cells (**Table S3**). Furthermore, V(D)J sequences from IgG^-^ DZ cells were enriched for canonical debilitating mutations, including stop codons and insertions or deletions that cause frameshift, and mutations affecting conserved residues (**Fig. 7B**).

**Figure 7.**
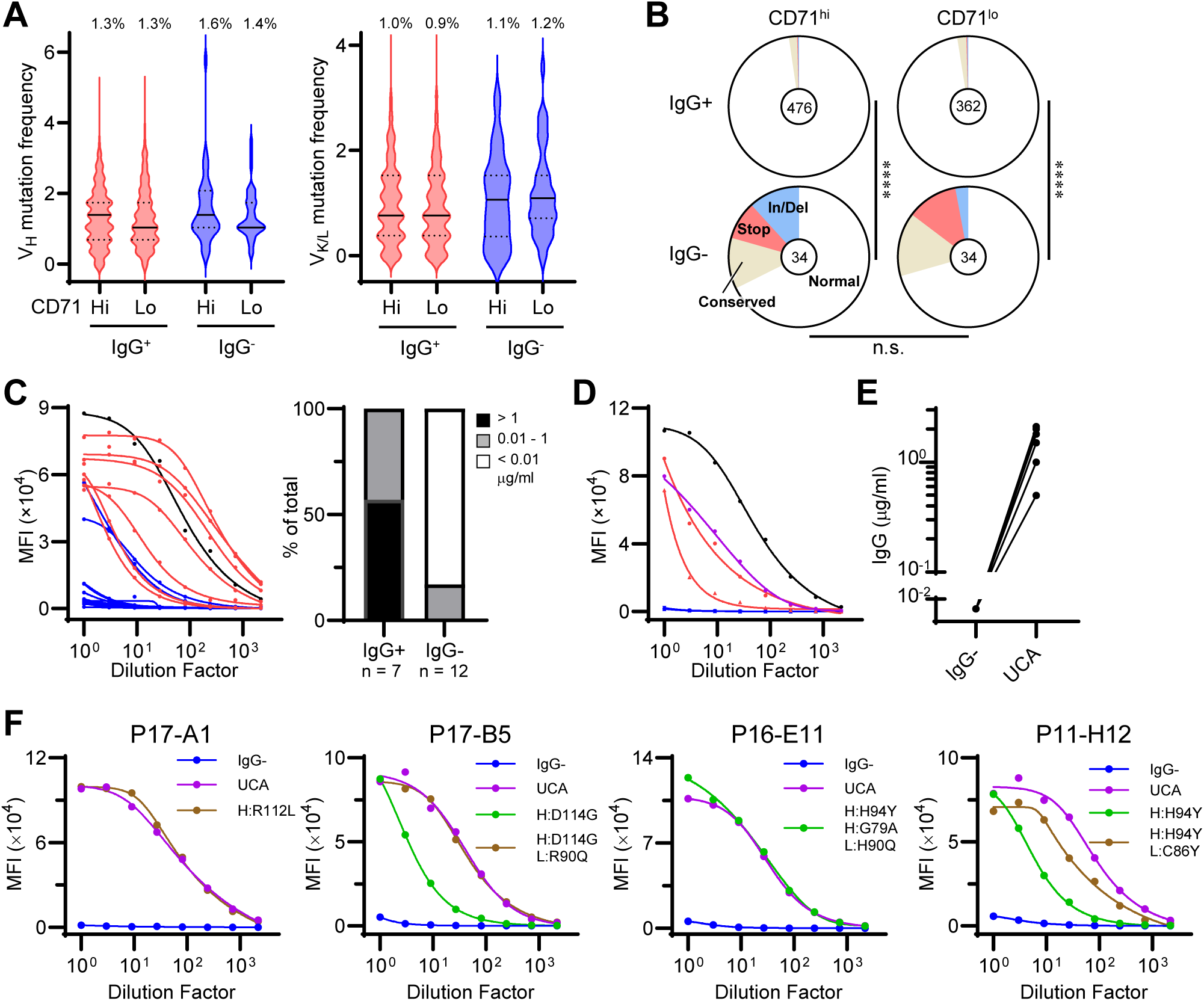
Hidden debilitating V(D)J mutations impair antibody expression. Paired Ig(H+L) V(D)J gene sequences were obtained from CD71^hi^ or CD71^lo^ DZ cells (see the legend of Fig. 6). (**A**) Violin plots show distributions of mutation frequency for V_H_ (left) and Vκ or Vλ (right) genes of indicated DZ subsets. Mutation frequencies represent number of nucleotide substitutions per base pairs sequenced for each V gene. No statistical differences were obtained among GC B cells by Kruskal-Walis test with Dunn’s multiple comparisons. (**B**) Pie charts show distributions of DZ cells expressing BCRs with the indicated classes of debilitating mutations: In/Del, insertions or deletions causing frameshifts; Stop, stop codons; Conserved, mutations affecting conserved immunoglobulin residues (see Materials and Methods); Normal, all remaining sequences. ****, *p* < 0.0001; n.s., *p* > 0.05 by Fisher’s exact test. (**C**) Recombinant antibody expression from V(D)J sequences of IgG^+^ (red, n = 7) and IgG^-^ (blue, n = 12) DZ subsets. Black, human 151K control IgG. Right panel shows percentage of recombinant antibodies at indicated IgG concentrations. (**D**) Recombinant antibody expression from V(D)J sequences of 2 IgG^+^ (red) and 3 IgG^-^ (blue) members of the same clonal lineage and their inferred UCA (purple). Black, 151K (human control IgG). (**E**) Recombinant antibody expression from V(D)J sequences of IgG^-^ DZ cells and their inferred UCA (n = 6 independent clonal lineages). (**F**) Recombinant antibody expression of IgG^-^ DZ cells and their UCA and reversion mutants.

Approximately two-thirds of paired V(D)J sequences from IgG^-^ DZ cells did not carry canonical debilitating mutations but appeared intact. To determine whether these apparently intact V(D)J sequences impaired antibody expression, we cloned paired Ig(H+L) V(D)J sequences from IgG^-^ DZ cells into standard IgG expression vectors and expressed as recombinant IgG in HEK293 cells. V(D)J sequences from IgG^+^ DZ cells efficiently produced recombinant IgG, whereas most V(D)J sequences from IgG^-^ DZ cells produced little or no IgG (**Fig. 7C**).

To determine whether these defects arose from SHM, we generated computationally inferred unmutated common ancestors (UCAs) from clonal lineages that contained both IgG^+^ and IgG^-^ members and compared recombinant IgG expression. Consistent with their original IgG secretion phenotypes, V(D)J sequences from IgG^+^ members efficiently produced recombinant IgG, while those from IgG^-^ members produced little or no recombinant IgG (**Fig. 7D**). For all cases we tested, V(D)J rearrangements from inferred UCAs efficiently produced recombinant IgG (**Figs. 7D** and **7E**). These results demonstrate that SHM produces both canonical and non-canonical debilitating mutations that impair IgG expression during GC responses.

We developed a logistic regression classifier trained on IgG^+^ and IgG^-^ antibody amino acid sequences (**Fig. S2**; **Methods**). This model allowed us to identify candidate amino acid substitutions predicted to impair antibody stability and/or protein expression. To test these predictions, we reverted candidate mutations to their corresponding UCA residues and assessed whether recombinant IgG expression could be restored. Mutations within CDR3 junctions were excluded from the analysis because it was not possible to determine whether they were introduced by AID-mediated SHM or during V(D)J recombination. Reversion of predicted destabilizing mutations either partially or completely restored recombinant IgG expression in all tested clonal lineages (**Fig. 7F**). The IgG^-^ clone P17-A1 carried 4 and 2 amino acid substitutions in the IgH and IgL V(D)J sequences, respectively. Reversion of the predicted unfavorable R112L heavy chain mutation restored recombinant IgG production to levels comparable to those of the UCA (**Fig. 7F**). Similarly, reversion of two predicted unfavorable mutations in the P17-B5 and three predicted unfavorable mutations in the P16-E11 clone restored recombinant IgG production to levels comparable to those of their respective UCAs (**Fig. 7F**). In the P17-B5 and P11-H12 IgG^-^ clones, we also observed an additive effect in which reversion of single unfavorable mutations partially restored recombinant IgG production, whereas combining reversion of a second unfavorable mutation either fully restored or substantially increased recombinant IgG levels (**Fig. 7F**). These results validate the predicted unfavorable mutations as functionally debilitating SHM products. Remarkably, in one case, complete restoration of IgG expression required reversion of only a single amino acid substitution. Together, these results indicate that productive IgG expression can be disrupted by surprisingly subtle changes in V(D)J sequences during SHM.

## Discussion

GC reactions dynamically generate clonal variants of B cells through SHM. Although apoptosis has been proposed to eliminate B cells expressing BCRs with lower-affinity and debilitating mutations, our findings support a broader role for apoptosis in regulating the persistence and composition of evolving GC populations. Ablation of BAK/BAX-dependent mitochondrial apoptosis elicited markedly enlarged and persistent GC responses that support B cell affinity maturation. Enlarged GC responses, however, accumulate atypical non-proliferating DZ populations that are normally constrained by mitochondrial apoptosis. Our findings suggest that mitochondrial apoptosis functions primarily as a quality-control mechanism that constrains the persistence of heterogeneous GC intermediates, thereby maintaining GC homeostasis.

Enlarged GC responses in BAKBAX^B-null^ mice efficiently support affinity maturation of B cells. This conclusion is based on direct measurement of antigen-binding avidity (as AvIn) of IgG produced by clonally expanded individual GC B cells rather than inference from enrichment of affinity-enhancing mutations ^47–50^. Importantly, enlarged GCs in BAKBAX^B-null^ mice contained antigen-specific B cells at frequencies comparable to or greater than those in B6 mice. Therefore, GC affinity maturation in BAKBAX^B-null^ mice cannot be explained by competition among only a small subset of antigen-specific B cells. Instead, affinity maturation is maintained in enlarged GC responses that contain large numbers of antigen-specific B cells. Our observations suggest that apoptosis is not required for selecting higher-affinity clones during GC responses. Rather, interaction with FDCs and competition for T cell help may be sufficient to drive GC affinity maturation ^51^. These interpretations are consistent with “permissive” selection in GCs, in which late GCs retain substantial clonal diversity and contain B cells spanning a broad range of affinities^25–27,52^.

A major finding of this study is the identification of non-proliferating Ki-67^lo^CD71^lo^ DZ populations that accumulated in BAKBAX^B-null^ mice but were rare in B6 mice. These non-proliferating DZ cells resemble the “seemingly quiescent” DZ populations reported in BCL-2 transgenic mice ^35^. Whether these “seemingly quiescent” DZ cells retain functional competence and how apoptosis regulates the persistence of these cells have not been extensively studied. CD71^lo^ DZ cells identified in this study retained capacity to proliferate, differentiate into plasmacytes, and secrete antibodies. Both CD71^lo^ DZ cells and CD71^hi^ DZ cells acquired comparable frequencies of V_H_ and V_K/L_ mutations, indicating that CD71^lo^ DZ cells are not long-lived quiescent cells persisting in GCs, but both DZ cell populations are similarly engaged in GC reactions. CD71^lo^ DZ cells frequently shared clonal lineages with CD71^hi^ DZ cells and were distributed throughout the clonal tree instead of confined to particular branches. We conclude that non-proliferating DZ cells represent transient GC intermediates that recurrently emerge from actively cycling GC populations.

Some clonal lineages were dominated by CD71^lo^ non-proliferating DZ B cells, whose BCR avidities (as AvIn) were higher than the median avidity of CD71^hi^ proliferating DZ B cells. Because GC affinity maturation is generally thought to progressively enrich higher-avidity B cell clones, we were surprised to find that these high-avidity lineages were dominated by CD71^lo^ non-proliferating DZ B cells, a cell compartment associated with clonal contraction during GC responses. We conclude that higher-avidity lineages can be extinguished from ongoing GC responses, and BCR avidity alone is not sufficient to predict the evolutionary trajectories of B cell lineages.

CD71^lo^ non-proliferating DZ states may represent transient cellular intermediates that arise from diverse cellular stresses generated during clonal selection and SHM in GCs. For example, BCR dysfunction resulting from debilitating V(D)J mutations could induce cell-cycle arrest and apoptosis. Other potential mechanisms include replication stress, which can induce cell-cycle arrest and apoptosis in GC B cells ^53^, DNA damage ^54^, metabolic insufficiency ^24^, off-target mutations ^55^, and other cellular stresses associated with extensive cell proliferation and high rates of mutation ^56,57^. Although speculative, these mechanisms are plausible as GCs are highly proliferative and mutagenic environment. Under physiological conditions, mitochondrial apoptosis may remove these highly stressed, transient intermediates before they accumulate and become detectable in wild-type GCs.

In contrast to the marked accumulation of non-proliferating DZ populations, LZ cells in BAKBAX^B-null^ mice appeared normal. One possible explanation is that apoptosis-resistant BAK/BAX-deficient LZ cells continue to reenter DZ program ^58^ and accumulate as non-proliferating DZ populations. Alternatively, additional death pathways independent of BAK/BAX-dependent mitochondrial apoptosis may contribute preferentially to purge LZ cells. In support of this possibility, “Rogue” GC B cell population including those that have acquired autoreactivity can be removed by the extrinsic pathway via the Fas:FasL interactions ^59^. GC B cells can also undergo necrosis in the absence of Caspase-9 ^60^, suggesting that compensatory non-apoptotic death pathways may also play a role in GC homeostasis when intrinsic apoptosis is disrupted.

Another unexpected feature of BAKBAX^B-null^ GC responses is the persistence of dysfunctional GC B cells carrying debilitating V(D)J mutations. Indeed, a subset of DZ B cells from BAKBAX^B-null^ mice expanded in Nojima cultures without secreting IgG. Both CD71^hi^ and CD71^lo^ DZ B cells generated IgG^-^ cultures, while GC B cells from B6 mice rarely did so. These IgG^-^ DZ B cells frequently carried canonical or non-canonical debilitating V(D)J mutations that impair antibody expression. It is likely that CD71^hi^ DZ B cells acquire debilitating mutations during proliferation and SHM. Accordingly, CD71^hi^IgG^-^ DZ B cells may represent recently generated dysfunctional intermediates, while CD71^lo^IgG^-^ DZ B cells may represent their non-proliferating descendants. That reversion of a single amino acid substitution often restored productive antibody expression indicates that antibody expression can easily be disrupted by SHM. Our findings extend a previous study that reported “structurally compromised” BCRs in DZ cells ^34^ by identifying causative, SHM-derived amino acid substitutions and showing that their reversion restores productive antibody expression.

Recent studies have shown that GC B cells transiently reduce SHM during clonal expansion ^61,62^. Whereas these studies define upstream mechanisms that limit ongoing mutational damage and preserve previously acquired affinity during rapid proliferation, our study identifies a distinct layer of cellular quality control revealed when mitochondrial apoptosis is disabled. In the absence of mitochondrial apoptosis, GCs accumulate cells in a non-proliferating state, cells belonging to lineages that are losing competitive fitness, and cells whose BCRs can no longer be productively expressed. We cannot distinguish whether apoptosis eliminates cells after they enter these states, or whether these states arise because cells that would otherwise have been eliminated persist. In either case, mitochondrial apoptosis constrains the persistence of GC B cells whose competence is compromised, a state not predicted by BCR avidity alone. Thus, SHM-induced loss of antibody expression represents one manifestation within a broader spectrum of cryptic GC states constrained by mitochondrial apoptosis.

Our findings support a revised model of GC homeostasis in which mitochondrial apoptosis is not required to maintain affinity maturation but acts as a cellular quality-control mechanism that constrains the size and cellular composition of evolving GC populations. Loss of BAK and BAX in B cells resulted in enlarged and prolonged GCs that contained heterogeneous non-proliferating DZ populations as well as variants with defective antibody expression. Affinity maturation nevertheless remained intact, demonstrating that it can be uncoupled from GC population dynamics. These cryptic GC B cell states reveal an underappreciated role for cellular competence in shaping clonal fitness during GC evolution.

### Limitation of the study

We demonstrated that CD71^lo^ DZ cells retained the capacity to proliferate, differentiate into plasmacytes, and secrete antibodies *ex vivo*. Whether these cells represent terminally arrested populations or re-enter cellular responses *in vivo* is unclear. Further studies will be required to determine their cellular fate and biological functions *in vivo* (*e.g.*, re-entry into the GC cycling program, differentiation into memory B cells or plasmacytes). We analyzed CD71^lo^ DZ cells that proliferated in Nojima cultures. This approach revealed both functionally competent CD71^lo^ DZ cells and cells carrying debilitating V(D)J mutations that impaired antibody expression. However, CD71^lo^ DZ cells that had irreversibly lost proliferative potential or were committed to cell death would not be sampled by our approach. Our analyses may underestimate the heterogeneity of non-proliferating DZ cells that accumulate in BAKBAX^B-null^ mice. Additional approaches that do not rely on cell proliferation *ex vivo* will help understand the full spectrum of non-proliferating DZ populations.

## Materials and Methods

### Mice and Immunizations

C57BL/6 mice and *Bax^tm2Sjk^Bak1^tm1Thsn^*/J (*Bak*^-/-^*Bax*^fl/fl^) mice were purchased from Jackson laboratory. *Bak*^-/-^*Bax*^fl/fl^ mice were backcrossed with B6 mice at least ten generations before experimental use. *Cd19*^Cre/wt^*Bak*^-/-^*Bax*^fl/fl^ mice (BAKBAX^B-null^ mice) were generated as previously described ^40^. All mice were maintained under specific pathogen-free conditions at the Duke University Animal Care Facility. Eight to 12-week-old female mice were immunized with 20 μg of recombinant protective antigens (rPA) from *Bacillus Anthracis* (BEI Resources) in the footpad of the right hind leg ^25^. Mice were analyzed 8 to 40 days after immunization. rPA antigens were mixed with Alhydrogel® adjuvant 2% (final 1%) before immunizations. All experiments involving animals were approved by the Duke University Institutional Animal Care and Use Committee.

### Flow Cytometry

GC B cells (GL-7^+^B220^hi^CD38^lo^IgD^-^CD138^-^) or LZ B cells (GL-7^+^B220^hi^CD38^lo^IgD^-^CXCR4^-^CD86^hi^) or DZ B cells (GL-7^+^B220^hi^CD38^lo^IgD^-^CXCR4^+^CD86^lo^) in the draining LNs were identified as described ^14,25^. In some experiments, DZ cells were further divided by surface expressions of CD71 or BCRs (Igκ and Igλ). For intracellular Ki-67 staining, samples were fixed after surface molecule labeling, permeabilized, and stained with Ki-67 antibody using BD Transcription Factor Buffer Set as described ^63^. Labeled cells were analyzed/sorted in a FACS Canto (BD Biosciences) or a FACS LSRII (BD Biosciences) or a FACS Vantage or a FACS Symphony S6 with DIVA option (BD Biosciences). Flow cytometric data were analyzed with FlowJo software (BD Biosciences). Doublets were excluded by FSC-A/FSC-H gating strategy. Cells that take up propidium iodide or labeled with LIVE/DEAD Fixable Near-IR Dead Cell Stain Kit (BD Biosciences) were excluded from our analysis. mAbs used in this study: FITC-or PE-conjugated anti-GL-7 (GL7, BD Biosciences or BioLegend, respectively), BV421-or BV785-conjugated anti-B220 (RA3-6B2, BioLegend), PE-Cy7-conjugated anti-CD38 (90, BioLegend), BV510-conjugated anti-IgD (11-26c.2a, BioLegend), PE-or APC-or BV605-conjugated anti-CD138 (281-2, BD Bioscience or BioLegend, respectively). PE-conjugated anti-CXCR4 (2B11/CXCR4, BD Biosciences), APC-conjugated anti-CD86 (GL-1, BioLegend), BV421-conjugated or biotinylated anti-CD71 (C2, BD Biosciences), BV421-or BV750-conjugated anti-Igκ (187.1, BD Biosciences), BV421-or BV750-conjugated anti-Igλ (R26-46, BD Biosciences), BV650-conjugated anti-Ki-67(B56, BD Biosciences), PC-CF594-conjugated anti-Bcl-6 (K112-91, BD Biosciences), PE-Cy5-conjugated anti-IgM (II/41, BD Biosciences), Biotinylated anti-CD95 (Jo2, BD Biosciences) and BV711-conjugated anti-MHCII (M5/114.15.2, BD Biosciences).

To compare the heterogeneity and subpopulation distribution of B cells between B6 and BAKBAX^B-null^ mice, high-dimensional flow cytometry data were analyzed using FlowJo software (BD Biosciences). Target B cell populations were first defined using standard manual gating strategies, and events within the B220^+^ B cell gates from both groups were then concatenated into a single data file to ensure an objective comparison within the exact same computational space. Unsupervised, non-linear dimensionality reduction was subsequently performed on the concatenated data using the Uniform Manifold Approximation and Projection (UMAP) plugin for FlowJo. The UMAP algorithm evaluated all markers to project high-dimensional single-cell features onto a two-dimensional plot (parameters were configured using Euclidean distance, with nearest neighbors set to 15 and minimum distance set to 0.5). Following UMAP visualization, specific germinal center B cell subpopulations, such as the LZ and DZ subsets, were identified based on their distinct marker expression profiles. Finally, the concatenated UMAP plot was de-concatenated by Sample ID to separate the data back into B6 and BAKBAX^B-null^ groups, facilitating the comparative analysis of population shifts and the identification of unique phenotypic accumulations selectively present in the BAK/BAX-deficient background.

### Single-cell Nojima culture

Sorted GC B cells were expanded in Nojima cultures as described previously ^25,44^. Briefly, NB-21.2D9 feeder cells were seeded into 96-well plates at 2,000 cells/well in B cell media (BCM); RPMI-1640 (Invitrogen) supplemented with 10% HyClone FBS (Thermo scientific), 5.5 × 10^-5^ M 2-mercaptoethanol, 10 mM HEPES, 1 mM sodium pyruvate, 100 units/ml penicillin, 100 μg/ml streptomycin, and MEM nonessential amino acid (all Invitrogen). Next day, recombinant mouse IL-4 (Peprotech; 2 ng/ml) was added to the cultures, and then single B cells were directly sorted into each well of 96-well plates using a FACS Vantage or a FACS Symphony S6. Two days after culture, 50% (vol.) of culture media were removed from cultures and 100% (vol.) of fresh BCM were added to the cultures. Two-thirds of the culture media were replaced with fresh BCM every day from day 4 to day 8. Culture supernatants were harvested on day 10. Culture plates were stored at -80°C until use for V(D)J amplifications.

### ELISA and Luminex assays

The presence of total and antigen-specific IgG in culture supernatants was determined by ELISA and Luminex multiplex assay ^25,44^. Diluted culture supernatants (1: 10 in PBS containing 0.5% BSA and 0.1% Tween-20) were screened for the presence of IgGs by standard ELISA. IgG^+^ samples were then screened for binding to rPA by Luminex assay. Briefly, culture supernatants were diluted (1: 10 or 1: 100) in 1×PBS containing 1% BSA, 0.05% NaN3 and 0.05% Tween20 (assay buffer) with 1% milk and incubated for 2 hours with the mixture of antigen-coupled microsphere beads in 96-well filter bottom plates (Millipore). After washing with assay buffer, these beads were incubated for 1 hour with PE-conjugated goat anti-mouse IgG Abs (Southern Biotech). After three washes, the beads were re-suspended in assay buffer, and the plates were read on a Bio-Plex 3D Suspension Array System (Bio-Rad). The following antigens were coupled with carboxylated beads (Luminex Corp): BSA (Affymetrix), goat anti-mouse Igκ, goat anti-mouse Igλ, goat anti-mouse IgG (all Southern Biotech), and rPA. For each IgG^+^ culture supernatant sample, concentrations of total IgG and rPA-specific IgG were determined in reference to monoclonal standard, BAP0105 (Abcam). AvIn values were obtained for rPA binding IgG samples in reference to BAP0105 ^25^.

### Amplification and analysis of V(D)J rearrangements

V(D)J rearrangements of cultured B cells were amplified by a semi-nested PCR as described previously ^44^. Total RNA was extracted from selected samples using Quick-RNA 96 kit (Zymo Research). cDNA was synthesized from DNase I-treated RNA using SMARTScribeTM Reverse Transcriptase (Clontech) with 0.2 μM each of gene-specific reverse primers (mIgG-RV1, mIgK-RV1, mIgL23-RV1, and mIgL145-RV1) and 1 μM of 5’ SMART template-switching oligo that contained plate-associated barcodes at 50°C for 50 min followed by 85°C for 5 min. cDNA was then subjected to two rounds of PCR using Herculase II fusion DNA polymerase (Agilent Technologies) with combinations of forward primers and reverse primers that contained well-associated barcodes (**Table S4**, RT-PCR A). Primary PCR: 95°C for 4 min, followed by 2 cycles of 95°C for 30 sec, 64°C for 20 sec, 72°C for 30 sec; 3 cycles of 95°C for 30 sec, 62°C for 20 sec, 72°C for 30 sec; 25 cycles of 95°C for 30 sec, 55°C for 20 sec, 72°C for 30 sec; and 72°C for 10 min. Secondary PCR: 95°C for 4 min, followed by 40 cycles of 95°C for 30 sec, 45°C for 20 sec, 72°C for 30 sec; and 72°C for 10 min.

In some experiments, cDNA was synthesized from DNase I-treated RNA using SuperScript III Reverse Transcriptase (Invitrogen) with 0.2 μM each of gene-specific reverse primers (mIgG-RV1, mIgKC2-RV1, mIgL23-RT, mIgLC1-RT, and mIgLC4-RT) at 50°C for 50 min followed by 85°C for 5 min. cDNA was then subjected to two rounds of PCR using Herculase II fusion DNA polymerase with combinations of forward primers and reverse primers that contained well-associated barcodes (**Table S4**, RT-PCR B). Primary PCR: 95°C for 4 min, followed by 6 cycles of 95°C for 30 sec, 62-52°C for 30 sec (-2°C per cycle), 72°C for 30 sec; 25 cycles of 95°C for 30 sec, 52°C for 30 sec, 72°C for 30 sec; and 72°C for 10 min. Secondary PCR: 95°C for 4 min, followed by 2 cycles of 95°C for 30 sec, 64°C for 20 sec, 72°C for 30 sec; 3 cycles of 95°C for 30 sec, 62°C for 20 sec, 72°C for 30 sec; 25 cycles of 95°C for 30 sec, 55°C for 20 sec, 72°C for 30 sec; and 72°C for 10 min.

Gel-purified, pooled V(D)J amplicands were submitted to DNA Link, Inc. to obtain DNA sequences using PacBio Sequel platform. Obtained circular consensus sequences (CCS) were analyzed to build a consensus sequence for each sample using AbSolute developed in the Kelsoe laboratory ^64^. The rearranged V, D, and J gene segments were first identified using IMGT/V-QUEST (http://www.imgt.org/) or Cloanalyst ^46^, and then numbers and kinds of point mutations were determined. Debilitating mutations included insertions or deletions causing frameshifts, stop codons, and mutations affecting conserved immunoglobulin residues critical for antibody structure and stability ^65^.

### Recombinant antibody expression and purification

Heavy-and light-chain variable domains of selected BCRs were cloned into human IgG1 and Igκ expression vectors (gift from Hedda Wardemann) ^66^. Recombinant antibodies were produced by transient transfection of Expi293F cells (according to manufacturer’s instruction) and purified from the culture supernatants using NAb protein G spin columns (Thermo Scientific). IgG concentrations were determined in reference to human myeloma protein 151K (Southern Biotech).

### Structural energetic modeling and prediction of IgG expression

For each antibody, mature heavy-and light-chain variable-region sequences were compared with their inferred unmutated common ancestor (UCA) sequences to identify amino acid substitutions acquired during SHM. Substitutions were defined in two directions: forward mutations from the UCA to the mature sequence, and reversion mutations from the mature sequence back to the corresponding UCA residue. Residue numbering was standardized using IMGT numbering implemented through ANARCI to ensure consistent mapping of substitutions across antibodies and between heavy and light chains ^67^.

Paired heavy-and light-chain variable-domain structures were predicted using IgFold ^68^. These IgFold-derived structures were used for all downstream energetic calculations to maintain a consistent structural representation across IgG^+^ and IgG^-^ antibodies. Before mutation modeling, each predicted structure was processed with FoldX RepairPDB to optimize side-chain conformations and reduce steric clashes. The total folding free energy of the repaired mature structure was extracted from the RepairPDB output and recorded as Stability dG (**Fig. S2C**) ^69^.

Per-mutation energetic effects were calculated with FoldX BuildModel. For each antibody, individual point substitutions were modeled in both the forward direction, representing mutations accumulated during affinity maturation (UCA → mature), and the reversion direction, representing restoration of mature residues to their inferred UCA state (mature → UCA). Mutation energies were extracted from FoldX BuildModel output files and used to construct two cumulative energetic features. Total_Positive_Energy was calculated as the sum of positive ΔΔG values from forward mutations and represents the cumulative destabilizing burden associated with somatic mutations (**Fig. S2A**). Total_Negative_Energy was calculated as the sum of negative ΔΔG values from reversion mutations and represents the cumulative stabilizing potential of restoring mature residues to their UCA state (**Fig. S2B**). Together with Stability_dG, these features provided a three-dimensional energetic representation of each mature antibody.

Before model training, antibodies containing classically defined debilitating mutations (*e.g.*, stop codons, changes in invariant residues, *i.e.*, Cysteines, Tryptophan) were excluded so that the model would focus on non-canonical energetic defects. IgG expression status was then modeled using logistic regression based on the three energetic features. Features were standardized by z-score transformation prior to training. Data were divided into stratified training and test sets (70% and 30%, respectively). To account for class imbalance in the training set, SMOTE (Synthetic Minority Over-sampling Technique) was applied within the training pipeline, and logistic regression was fit using balanced class weights and L2 regularization. The regularization parameter was selected by stratified five-fold cross-validation using average precision as the optimization metric. Final model performance was evaluated on the held-out test set.

For model-guided reversion design, candidate mutations were restricted to amino acid substitutions present in the mature antibody and therefore eligible for reversion to the corresponding UCA residue. Individual amino acid reversions were ranked according to their contributions to the Total_Negative_Energy, with larger stabilizing contributions prioritized. Reversion variants were generated iteratively by progressively adding top-ranked reversions and recalculating the predicted probability of IgG expression using the trained logistic regression model. Candidate variants containing a minimal number of reversions predicted to be IgG^+^ were selected for experimental testing of recombinant IgG expression.

### Statistics

Statistical significance (*p* < 0.05) was determined by Two-way Mixed-effects model, Wilcoxon matched-pairs signed rank test, Kruskal-Walis test with Dunn’s multiple comparisons, Unpaired *t* test with Welch’s correction, One-tailed Spearman’s correlation test, Mann-Whitney’s U test, and Fisher’s exact test using GraphPad Prism software (version 11.0.0, GraphPad Software). Statistic tests are indicated within each figure legend.

## Acknowledgments

We thank J. Finney, X. Nie, B. Pun, P. Liu, C. Nishihara, and O. Balmert for assistance. We thank all members of Kuraoka, Kelsoe, and Wiehe labs for support. We thank S. Harrison for discussion and comments on this study. We thank H. Wardemann for reagents. We thank B. Herold and M. Gromisch for LN samples from ΔgD-2 immunized mice. This work was supported in part by grants from the US NIH UM1 AI144371 (G.K.) and R01 AI177673 (M.K.).

## Author contributions

C-H.Y., A.W., D.L., X.L., W.Z. and M.K. performed experiments; S.S. developed V(D)J sequencing method; N.S., H.K. and K.W. developed mathematical modeling; C-H.Y., N.S., H.K., A.W., D.L., X.L., K.W., G.K. and M.K. analyzed data; K.W., G.K., and M.K. designed experiments; G.K. and M.K. conceptualized the study; C-H.Y. and M.K. wrote the paper; N.S., H.K., K.W., and G.K. edited the paper.

## Declaration of generative AI and AI-assisted technologies in the writing process

During the preparation of this work, the authors used ChatGPT (OpenAI) to improve the language and readability of the manuscript. After using this tool or service, the authors reviewed and edited the content as needed and take full responsibility for the publication’s content.

---

**Figure S1.**
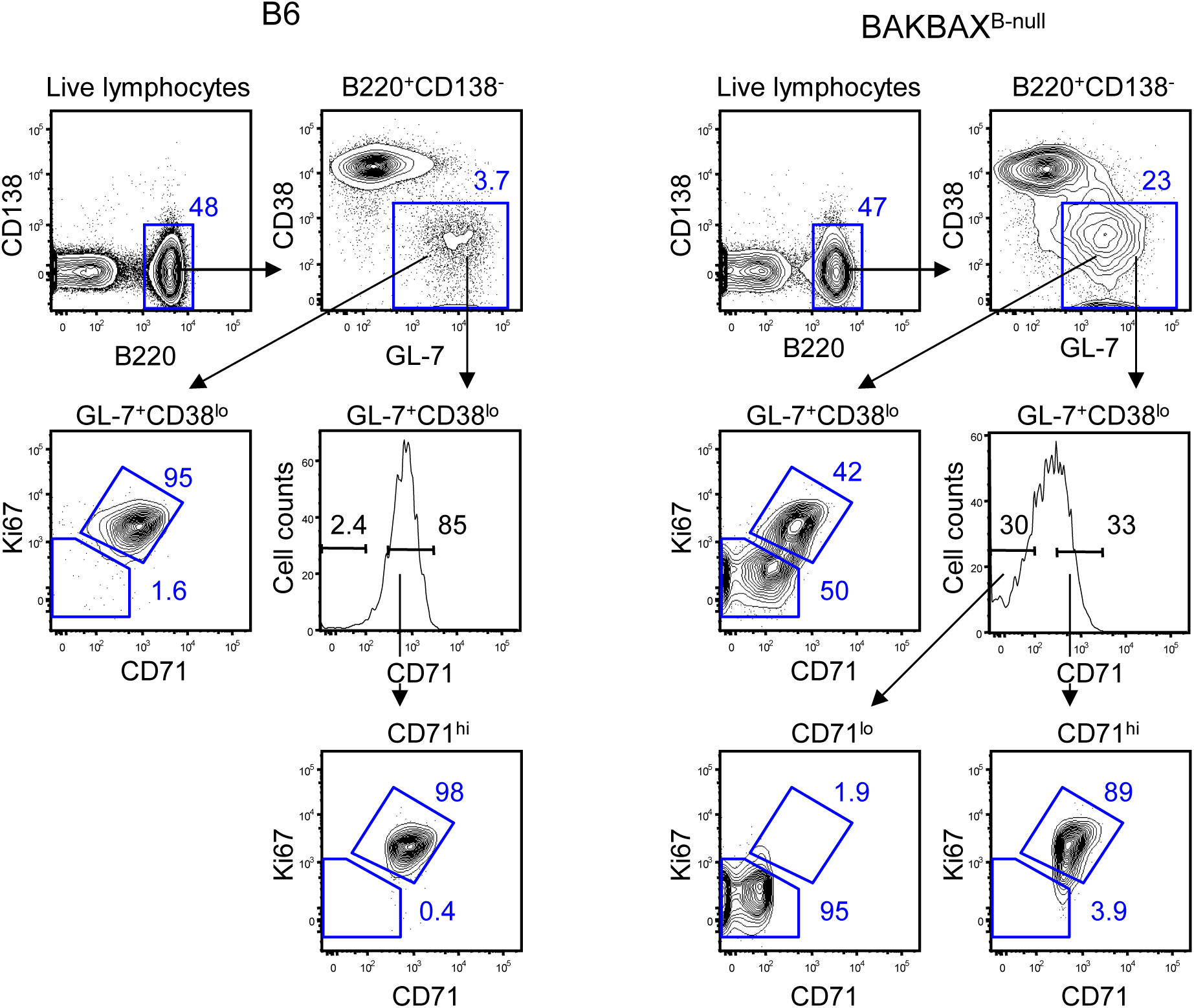
Association of intracellular Ki-67 and surface CD71 expressions in GC B cells. GC B cells (top panels) were further analyzed based on surface CD71 expression (x-axis) and intracellular Ki-67 expression (y-axis). Virtually all B6 GC B cells exhibited a Ki-67^hi^CD71^lo^ phenotype, whereas BAKBAX^B-null^ GC B cells contained substantial Ki-67^hi^CD71^hi^ and Ki-67^lo^CD71^lo^ populations (middle contour plots). Although CD71 expressions did not exhibit clearly separable peaks (middle histograms), gating the top 30% and bottom 30% of CD71-expressing cells successfully enriched Ki-67^hi^CD71^hi^ and Ki-67^lo^CD71^lo^ populations, respectively (bottom panels).

**Figure S2.**
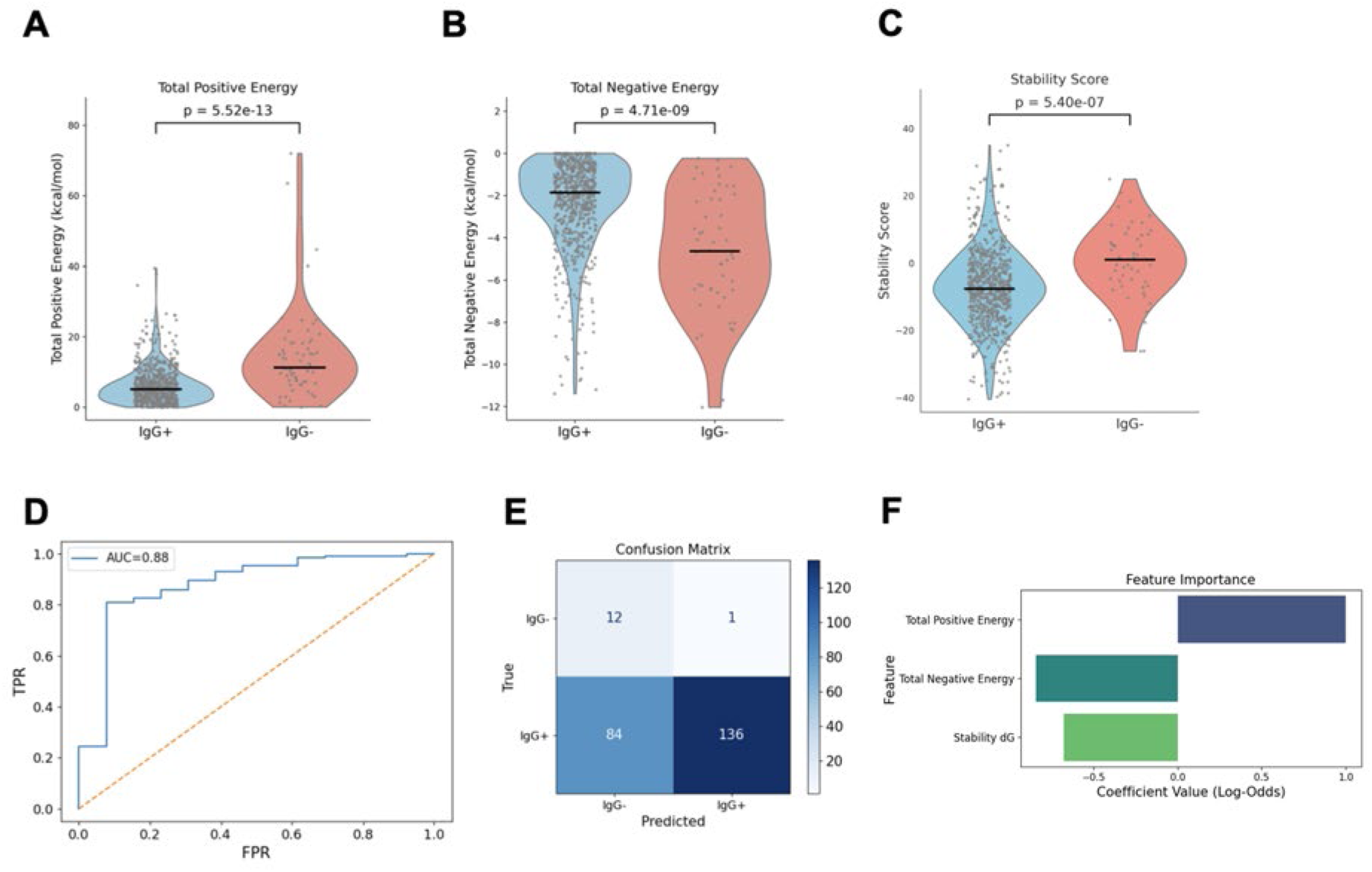
Modeling framework to predict IgG expression and identify candidate amino acid substitutions associated with impaired IgG expression. A logistic regression classifier was trained on features of structures predicted from antibody sequences of IgG^+^ and IgG^-^ DZ B cells. (**A** – **C**) Distributions of feature values for IgG^+^ vs. IgG^-^ sequences used in the model training. (**A**) Total positive energy, calculated as the sum of the FoldX estimated ΔΔG contributions of all predicted destabilizing mutations (*i.e.*, mutations with positive ΔΔG values) accumulated along the forward maturation pathway from the inferred unmutated common ancestor (UCA) to the mature antibody. Larger positive values indicate a greater cumulative destabilizing effect. (**B**) Total negative energy, calculated as the sum of the FoldX estimated ΔΔG contributions of all predicted stabilizing mutations (*i.e.*, mutations with negative ΔΔG values) accumulated along the reverse pathway from the mature antibody to the inferred UCA. More negative values indicate a greater cumulative stabilizing effect. (**C**) Stability score, calculated as the FoldX predicted structural stability dG of the mature antibody. More negative values indicate greater predicted stability. (**D**) Receiver operating characteristic (ROC) curve showing classifier performance. (**E**) Confusion matrix summarizing classification accuracy. (**F**) Feature-importance plot showing the logistic regression coefficient weights for the structural features included in the model.

**Table S1.** V_H_ gene segment usage of GC B-cell subsets.

|  | B6<br>GC d8<br>n = 469 | B6<br>GC d16<br>n = 179 | B6<br>GC d24<br>n = 277 | BAKBAX<br>GC d8<br>n = 164 | BAKBAX<br>GC d16<br>n = 229 | BAKBAX<br>GC d24<br>n = 494 | CD71<sup>hi</sup><br>DZ IgG<sup>+</sup><br>n = 476 | CD71<sup>lo</sup><br>DZ IgG<sup>+</sup><br>n = 362 | CD71<sup>hi</sup><br>DZ IgG<sup>-</sup><br>n = 34 | CD71<sup>lo</sup><br>DZ IgG<sup>-</sup><br>n = 34 |
| --- | --- | --- | --- | --- | --- | --- | --- | --- | --- | --- |
| 1-85 | 1.5 | 1.7 | 0.4 | 0.0 | 0.9 | 5.5 | 6.3 | 2.8 | 2.9 | 5.9 |
| 1-84 | 0.0 | 0.0 | 1.1 | 0.0 | 0.0 | 0.0 | 0.0 | 0.0 | 0.0 | 0.0 |
| 1-82 | 10.9 | 3.9 | 5.8 | 4.9 | 14.8 | 3.4 | 2.3 | 13.5 | 0.0 | 5.9 |
| 1-81 | 0.2 | 4.5 | 11.9 | 1.2 | 3.9 | 3.6 | 2.3 | 1.4 | 2.9 | 2.9 |
| 1-80 | 5.3 | 2.8 | 0.4 | 5.5 | 0.4 | 1.8 | 1.1 | 2.2 | 0.0 | 0.0 |
| 1-78 | 0.9 | 1.1 | 4.3 | 0.0 | 0.0 | 0.0 | 0.0 | 0.6 | 0.0 | 0.0 |
| 1-77 | 0.4 | 0.6 | 0.0 | 0.6 | 0.4 | 0.0 | 0.4 | 1.7 | 0.0 | 0.0 |
| 1-76 | 0.2 | 1.7 | 1.4 | 0.6 | 0.4 | 1.0 | 0.2 | 0.6 | 0.0 | 0.0 |
| 1-75 | 0.6 | 2.8 | 1.1 | 0.6 | 0.4 | 1.4 | 2.5 | 0.3 | 2.9 | 0.0 |
| 1-74 | 0.9 | 0.6 | 0.4 | 1.8 | 0.4 | 1.4 | 1.7 | 0.6 | 0.0 | 8.8 |
| 1-72 | 0.2 | 0.0 | 1.8 | 0.0 | 3.1 | 2.2 | 1.3 | 8.0 | 0.0 | 2.9 |
| 8-12 | 1.1 | 0.6 | 0.4 | 0.6 | 0.9 | 0.6 | 1.7 | 3.6 | 0.0 | 0.0 |
| 1-69 | 1.3 | 0.6 | 0.4 | 1.8 | 3.9 | 2.0 | 0.4 | 1.4 | 0.0 | 0.0 |
| 1-67 | 0.0 | 0.6 | 0.0 | 0.0 | 0.0 | 0.0 | 0.0 | 0.0 | 0.0 | 0.0 |
| 1-66 | 0.2 | 2.8 | 1.1 | 0.0 | 0.0 | 0.2 | 0.0 | 1.7 | 0.0 | 0.0 |
| 1-64 | 1.1 | 3.4 | 3.6 | 0.6 | 3.5 | 1.2 | 0.8 | 1.4 | 0.0 | 0.0 |
| 1-63 | 0.0 | 0.0 | 0.0 | 0.0 | 0.9 | 0.6 | 0.0 | 0.0 | 0.0 | 0.0 |
| 1-62-2/1-71 | 0.4 | 1.1 | 0.7 | 0.0 | 0.0 | 0.4 | 0.4 | 0.0 | 2.9 | 0.0 |
| 1-61 | 0.9 | 1.1 | 0.4 | 1.8 | 0.0 | 0.0 | 0.0 | 0.0 | 0.0 | 0.0 |
| 1-59 | 0.6 | 0.0 | 3.6 | 3.7 | 0.9 | 0.4 | 2.1 | 1.7 | 0.0 | 0.0 |
| 1-58 | 0.0 | 1.7 | 0.0 | 1.2 | 0.0 | 0.0 | 0.8 | 0.6 | 0.0 | 0.0 |
| 8-8 | 9.6 | 1.7 | 2.2 | 4.9 | 7.0 | 1.8 | 3.2 | 6.1 | 0.0 | 2.9 |
| 1-55 | 0.6 | 1.7 | 3.6 | 0.6 | 2.2 | 0.6 | 5.0 | 3.0 | 5.9 | 5.9 |
| 1-54 | 0.2 | 0.6 | 0.0 | 0.0 | 4.4 | 0.0 | 0.0 | 0.3 | 2.9 | 0.0 |
| 1-53 | 1.3 | 1.1 | 7.9 | 5.5 | 3.9 | 5.5 | 5.9 | 3.6 | 5.9 | 2.9 |
| 1-52 | 1.9 | 3.4 | 2.2 | 4.9 | 0.9 | 1.8 | 0.6 | 0.0 | 0.0 | 0.0 |
| 1-50 | 1.5 | 0.6 | 0.7 | 0.0 | 1.3 | 1.4 | 1.3 | 0.6 | 2.9 | 0.0 |
| 1-49 | 0.0 | 0.0 | 0.0 | 0.0 | 0.0 | 0.0 | 0.0 | 0.0 | 0.0 | 2.9 |
| 1-47 | 0.2 | 0.6 | 0.0 | 0.0 | 0.0 | 0.0 | 0.0 | 0.0 | 0.0 | 0.0 |
| 1-43 | 0.0 | 0.0 | 0.0 | 0.0 | 0.0 | 0.2 | 0.0 | 0.0 | 0.0 | 0.0 |
| 1-42 | 1.3 | 1.1 | 0.7 | 0.0 | 3.1 | 0.6 | 6.1 | 1.7 | 14.7 | 2.9 |
| 1-39 | 0.0 | 1.7 | 1.8 | 0.0 | 0.4 | 0.8 | 1.7 | 2.5 | 0.0 | 0.0 |
| 1-36 | 0.0 | 0.0 | 1.4 | 0.0 | 0.0 | 0.4 | 0.0 | 0.0 | 0.0 | 0.0 |
| 1-34 | 0.2 | 0.0 | 0.0 | 0.0 | 0.0 | 2.4 | 1.9 | 0.0 | 0.0 | 0.0 |
| 1-31 | 0.0 | 0.0 | 0.0 | 0.0 | 0.0 | 8.7 | 0.4 | 0.0 | 0.0 | 0.0 |
| 1-26 | 1.3 | 3.9 | 6.5 | 9.8 | 12.7 | 6.7 | 14.3 | 9.4 | 8.8 | 11.8 |
| 1-22 | 0.0 | 0.0 | 0.4 | 0.6 | 0.0 | 2.6 | 2.3 | 4.7 | 0.0 | 0.0 |
| 1-20 | 0.0 | 0.0 | 0.0 | 0.0 | 0.0 | 1.6 | 1.5 | 2.5 | 0.0 | 2.9 |
| 1-19 | 0.2 | 3.9 | 1.1 | 0.0 | 3.1 | 9.7 | 0.2 | 0.6 | 0.0 | 0.0 |
| 1-18 | 0.2 | 0.0 | 0.4 | 3.0 | 0.4 | 0.2 | 0.4 | 0.3 | 0.0 | 0.0 |
| 1-15 | 0.6 | 0.6 | 2.2 | 0.0 | 0.0 | 0.4 | 0.2 | 1.7 | 2.9 | 0.0 |
| 1-12 | 0.2 | 0.6 | 0.4 | 0.0 | 0.0 | 0.0 | 2.3 | 0.0 | 0.0 | 2.9 |
| 1-11 | 0.0 | 0.0 | 0.0 | 0.0 | 0.0 | 0.0 | 0.0 | 0.0 | 5.9 | 0.0 |
| 1-9 | 0.9 | 1.7 | 1.1 | 0.0 | 0.0 | 0.6 | 1.5 | 1.9 | 2.9 | 0.0 |
| 15-2 | 0.0 | 0.0 | 0.0 | 0.0 | 0.9 | 0.0 | 0.0 | 0.0 | 0.0 | 0.0 |
| 1-7 | 0.0 | 0.6 | 0.4 | 0.0 | 0.9 | 1.0 | 0.4 | 0.0 | 0.0 | 0.0 |
| 10-3 | 0.0 | 0.0 | 1.4 | 0.0 | 1.3 | 0.0 | 0.0 | 0.0 | 0.0 | 0.0 |
| 1-5 | 0.0 | 1.7 | 0.4 | 0.6 | 0.4 | 0.2 | 1.9 | 0.0 | 5.9 | 0.0 |
| 1-4 | 0.2 | 2.8 | 0.4 | 0.0 | 0.0 | 0.0 | 0.0 | 0.0 | 0.0 | 0.0 |
| 10-1 | 0.0 | 1.7 | 0.0 | 0.0 | 0.0 | 5.1 | 0.0 | 0.0 | 0.0 | 0.0 |
| 6-6 | 0.0 | 1.7 | 0.7 | 0.0 | 0.9 | 0.0 | 0.2 | 0.3 | 0.0 | 0.0 |
| 6-3 | 0.0 | 0.6 | 1.4 | 0.6 | 0.0 | 1.2 | 0.2 | 0.0 | 2.9 | 0.0 |
| 3-8 | 0.0 | 0.0 | 0.7 | 0.0 | 0.0 | 0.4 | 0.0 | 0.0 | 0.0 | 0.0 |
| 9-4 | 0.0 | 0.6 | 0.0 | 0.0 | 0.0 | 0.6 | 3.2 | 0.3 | 2.9 | 5.9 |
| 3-6 | 0.9 | 1.1 | 4.0 | 0.0 | 1.7 | 1.2 | 1.9 | 0.3 | 0.0 | 2.9 |
| 3-4 | 0.2 | 0.0 | 0.4 | 0.0 | 0.0 | 0.0 | 0.0 | 0.0 | 0.0 | 0.0 |
| 14-4 | 40.7 | 0.6 | 2.5 | 34.1 | 3.1 | 1.0 | 1.3 | 6.9 | 0.0 | 14.7 |
| 7-3 | 0.2 | 1.1 | 0.0 | 0.0 | 0.0 | 0.2 | 1.1 | 0.3 | 0.0 | 0.0 |
| 9-3 | 0.4 | 2.2 | 1.1 | 0.6 | 4.8 | 0.4 | 1.5 | 3.6 | 5.9 | 14.7 |
| 9-1 | 1.7 | 0.0 | 0.4 | 0.6 | 0.0 | 0.4 | 0.2 | 0.3 | 2.9 | 0.0 |
| 14-3 | 0.0 | 1.7 | 0.4 | 0.0 | 3.9 | 7.5 | 0.2 | 0.0 | 0.0 | 0.0 |
| 14-2 | 3.4 | 19.0 | 4.0 | 0.6 | 0.9 | 3.2 | 5.7 | 1.4 | 2.9 | 0.0 |
| 3-1 | 0.2 | 0.6 | 0.0 | 0.0 | 0.4 | 0.4 | 0.0 | 1.1 | 0.0 | 0.0 |
| 4-1 | 0.2 | 0.0 | 0.0 | 0.0 | 0.0 | 0.0 | 0.2 | 0.0 | 0.0 | 0.0 |
| 14-1 | 3.0 | 1.7 | 1.1 | 3.0 | 0.9 | 0.6 | 0.2 | 0.8 | 2.9 | 0.0 |
| 2-9 | 0.2 | 0.0 | 0.0 | 1.8 | 0.0 | 0.0 | 0.0 | 0.0 | 0.0 | 0.0 |
| 5-17 | 0.2 | 1.1 | 0.7 | 0.0 | 1.3 | 0.8 | 0.4 | 1.4 | 0.0 | 0.0 |
| 5-16 | 0.2 | 0.6 | 0.0 | 0.0 | 0.0 | 0.0 | 0.4 | 0.6 | 0.0 | 0.0 |
| 2-9-1 | 0.2 | 0.0 | 0.0 | 0.0 | 0.0 | 0.0 | 0.0 | 0.3 | 0.0 | 0.0 |
| 5-9-1 | 0.2 | 0.6 | 0.0 | 0.0 | 0.0 | 0.0 | 0.0 | 0.3 | 0.0 | 0.0 |
| 2-6 | 0.0 | 0.0 | 0.4 | 1.2 | 0.0 | 0.2 | 5.9 | 0.3 | 5.9 | 0.0 |
| 5-12 | 0.0 | 0.6 | 0.0 | 0.0 | 0.0 | 0.0 | 0.2 | 0.0 | 0.0 | 0.0 |
| 2-5 | 0.0 | 1.1 | 0.0 | 0.0 | 0.9 | 0.0 | 0.0 | 0.0 | 0.0 | 0.0 |
| 2-4 | 0.0 | 0.0 | 0.0 | 1.2 | 0.0 | 0.0 | 0.0 | 0.0 | 0.0 | 0.0 |
| 5-6 | 0.4 | 3.9 | 5.4 | 0.6 | 2.6 | 2.6 | 1.3 | 0.8 | 2.9 | 0.0 |
| 2-3 | 0.0 | 0.0 | 0.4 | 0.0 | 0.0 | 0.0 | 0.0 | 0.6 | 0.0 | 0.0 |
| 5-4 | 0.2 | 0.6 | 0.4 | 0.6 | 0.9 | 0.0 | 0.6 | 0.0 | 0.0 | 0.0 |
| 2-2 | 0.2 | 0.0 | 2.5 | 0.0 | 0.0 | 0.8 | 0.0 | 0.3 | 0.0 | 0.0 |

The percentages of V<sub>H</sub> gene segments among all V<sub>H</sub> gene segments within each B-cell subset are shown.

**Table S2.** Number of clonal cultures with or without IgG^1^.

| GC B cells | IgG<sup>+</sup> | IgG<sup>-</sup> | Ratio (IgG<sup>-</sup>/IgG<sup>+</sup>) |
| --- | --- | --- | --- |
| BAKBAX<sup>B-null</sup> CD71<sup>hi</sup> DZ<sup>2</sup> | 208 | 64 | 0.31 |
| BAKBAX<sup>B-null</sup> CD71<sup>lo</sup> DZ<sup>2</sup> | 146 | 51 | 0.35 |
| B6 GC DZ<sup>2</sup> | 54 | 1 | 0.02 |
| B6 GC<sup>3</sup> | 129 | 0 | 0.00 |

<sup>1</sup> On days 5-8 of Nojima cultures, presence of expanded single B cells was determined using light microscopy. After Nojima cultures, IgG secretion was determined by ELISA. IgG<sup>+</sup> represents clonal cultures that contain IgG. IgG<sup>-</sup> represents clonal cultures that contain expanded B cells without secreting IgG.
<sup>2</sup> DZ B cells were sorted from popliteal LNs of B6 or BAKBAX<sup>B-null</sup> mice following footpad immunization with rPA in alhydrogel.
<sup>3</sup> GC B cells were sorted from inguinal LNs of B6 mice following s.c. immunization with ΔgD-2 herpes virus vaccine candidate.

**Table S3.**
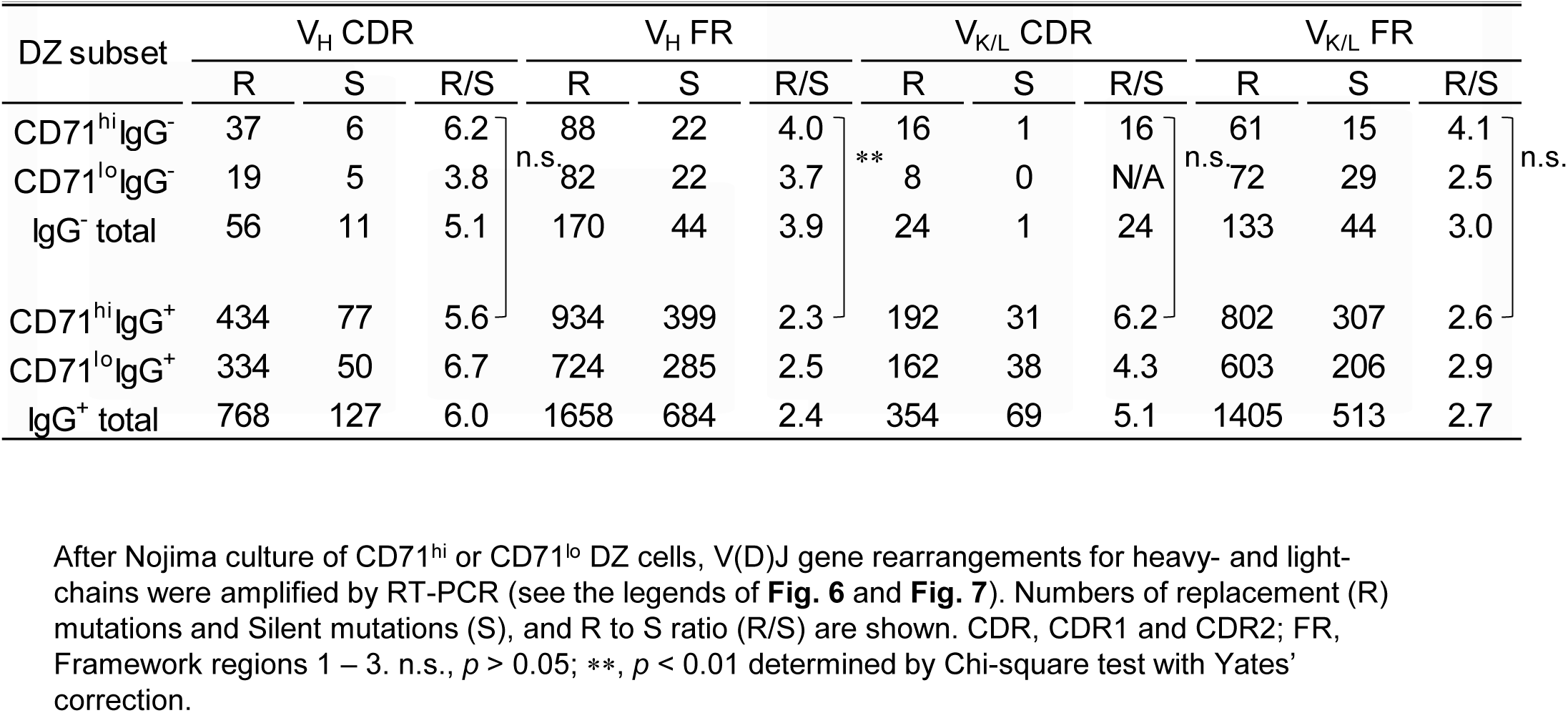
Number of Replacement and Silent mutations in V_H_ and V_K/L_ genes.

**Table S4.**
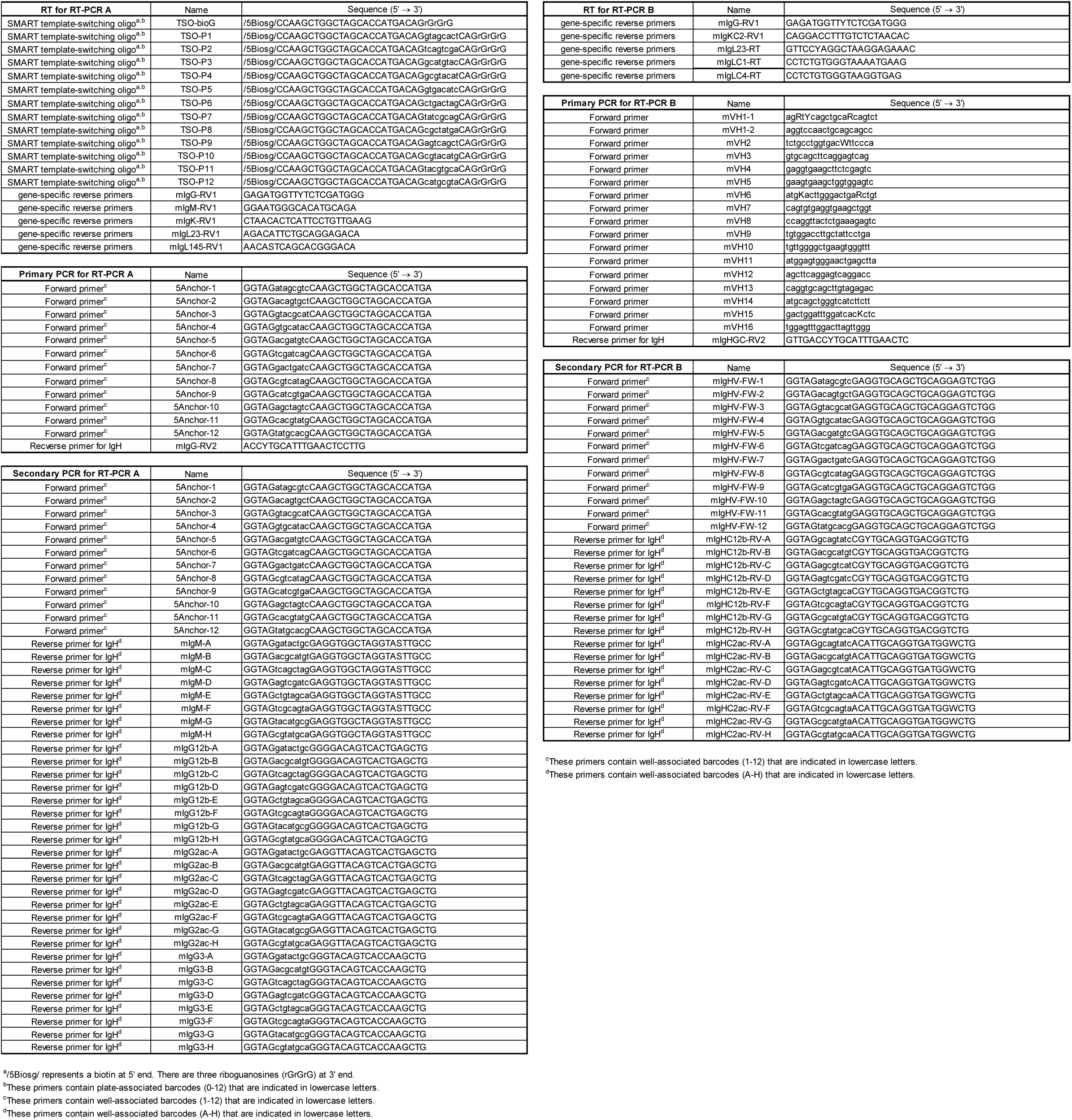
Primer sequences.

## References

1. Griffiths, G.M., Berek, C., Kaartinen, M., and Milstein, C. (1984). Somatic mutation and the maturation of immune response to 2-phenyl oxazolone. Nature 312, 271–275. 10.1038/312271a0.

2. Berek, C., Berger, A., and Apel, M. (1991). Maturation of the immune response in germinal centers. Cell 67, 1121–1129. 10.1016/0092-8674(91)90289-b.

3. Jacob, J., Kelsoe, G., Rajewsky, K., and Weiss, U. (1991). Intraclonal generation of antibody mutants in germinal centres. Nature 354, 389–392. 10.1038/354389a0.

4. MacLennan, I.C. (1994). Germinal centers. Annu Rev Immunol 12, 117–139. 10.1146/annurev.iy.12.040194.001001.

5. Rajewsky, K. (1996). Clonal selection and learning in the antibody system. Nature 381, 751–758. 10.1038/381751a0.

6. Mesin, L., Ersching, J., and Victora, G.D. (2016). Germinal Center B Cell Dynamics. Immunity 45, 471–482. 10.1016/j.immuni.2016.09.001.

7. Lau, A.W., and Brink, R. (2020). Selection in the germinal center. Curr Opin Immunol 63, 29–34. 10.1016/j.coi.2019.11.001.

8. Victora, G.D., and Nussenzweig, M.C. (2022). Germinal Centers. Annu Rev Immunol 40, 413–442. 10.1146/annurev-immunol-120419-022408.

9. Kepler, T.B., and Perelson, A.S. (1993). Cyclic re-entry of germinal center B cells and the efficiency of affinity maturation. Immunol Today 14, 412–415. 10.1016/0167-5699(93)90145-B.

10. Oprea, M., and Perelson, A.S. (1997). Somatic mutation leads to efficient affinity maturation when centrocytes recycle back to centroblasts. J Immunol 158, 5155–5162.

11. Allen, C.D., Okada, T., Tang, H.L., and Cyster, J.G. (2007). Imaging of germinal center selection events during affinity maturation. Science 315, 528–531. 10.1126/science.1136736.

12. Hauser, A.E., Junt, T., Mempel, T.R., Sneddon, M.W., Kleinstein, S.H., Henrickson, S.E., von Andrian, U.H., Shlomchik, M.J., and Haberman, A.M. (2007). Definition of germinal-center B cell migration in vivo reveals predominant intrazonal circulation patterns. Immunity 26, 655–667. 10.1016/j.immuni.2007.04.008.

13. Schwickert, T.A., Lindquist, R.L., Shakhar, G., Livshits, G., Skokos, D., Kosco-Vilbois, M.H., Dustin, M.L., and Nussenzweig, M.C. (2007). In vivo imaging of germinal centres reveals a dynamic open structure. Nature 446, 83–87. 10.1038/nature05573.

14. Victora, G.D., Schwickert, T.A., Fooksman, D.R., Kamphorst, A.O., Meyer-Hermann, M., Dustin, M.L., and Nussenzweig, M.C. (2010). Germinal center dynamics revealed by multiphoton microscopy with a photoactivatable fluorescent reporter. Cell 143, 592–605. 10.1016/j.cell.2010.10.032.

15. Allen, C.D., and Cyster, J.G. (2008). Follicular dendritic cell networks of primary follicles and germinal centers: phenotype and function. Semin Immunol 20, 14–25. 10.1016/j.smim.2007.12.001.

16. Crotty, S. (2019). T Follicular Helper Cell Biology: A Decade of Discovery and Diseases. Immunity 50, 1132–1148. 10.1016/j.immuni.2019.04.011.

17. Bannard, O., Horton, R.M., Allen, C.D., An, J., Nagasawa, T., and Cyster, J.G. (2013). Germinal center centroblasts transition to a centrocyte phenotype according to a timed program and depend on the dark zone for effective selection. Immunity 39, 912–924. 10.1016/j.immuni.2013.08.038.

18. Grootveld, A.K., Kyaw, W., Panova, V., Lau, A.W.Y., Ashwin, E., Seuzaret, G., Dhenni, R., Bhattacharyya, N.D., Khoo, W.H., Biro, M., et al. (2023). Apoptotic cell fragments locally activate tingible body macrophages in the germinal center. Cell 186, 1144–1161 e1118. 10.1016/j.cell.2023.02.004.

19. Mueller, J., Matloubian, M., and Zikherman, J. (2015). Cutting edge: An in vivo reporter reveals active B cell receptor signaling in the germinal center. J Immunol 194, 2993–2997. 10.4049/jimmunol.1403086.

20. Chen, S.T., Oliveira, T.Y., Gazumyan, A., Cipolla, M., and Nussenzweig, M.C. (2023). B cell receptor signaling in germinal centers prolongs survival and primes B cells for selection. Immunity 56, 547–561 e547. 10.1016/j.immuni.2023.02.003.

21. Dominguez-Sola, D., Victora, G.D., Ying, C.Y., Phan, R.T., Saito, M., Nussenzweig, M.C., and Dalla-Favera, R. (2012). The proto-oncogene MYC is required for selection in the germinal center and cyclic reentry. Nat Immunol 13, 1083–1091. 10.1038/ni.2428.

22. Gitlin, A.D., Shulman, Z., and Nussenzweig, M.C. (2014). Clonal selection in the germinal centre by regulated proliferation and hypermutation. Nature 509, 637–640. 10.1038/nature13300.

23. Shulman, Z., Gitlin, A.D., Weinstein, J.S., Lainez, B., Esplugues, E., Flavell, R.A., Craft, J.E., and Nussenzweig, M.C. (2014). Dynamic signaling by T follicular helper cells during germinal center B cell selection. Science 345, 1058–1062. 10.1126/science.1257861.

24. Ersching, J., Efeyan, A., Mesin, L., Jacobsen, J.T., Pasqual, G., Grabiner, B.C., Dominguez-Sola, D., Sabatini, D.M., and Victora, G.D. (2017). Germinal Center Selection and Affinity Maturation Require Dynamic Regulation of mTORC1 Kinase. Immunity 46, 1045–1058 e1046. 10.1016/j.immuni.2017.06.005.

25. Kuraoka, M., Schmidt, A.G., Nojima, T., Feng, F., Watanabe, A., Kitamura, D., Harrison, S.C., Kepler, T.B., and Kelsoe, G. (2016). Complex Antigens Drive Permissive Clonal Selection in Germinal Centers. Immunity 44, 542–552. 10.1016/j.immuni.2016.02.010.

26. Tas, J.M., Mesin, L., Pasqual, G., Targ, S., Jacobsen, J.T., Mano, Y.M., Chen, C.S., Weill, J.C., Reynaud, C.A., Browne, E.P., et al. (2016). Visualizing antibody affinity maturation in germinal centers. Science 351, 1048–1054. 10.1126/science.aad3439.

27. Sprumont, A., Rodrigues, A., McGowan, S.J., Bannard, C., and Bannard, O. (2023). Germinal centers output clonally diverse plasma cell populations expressing high-and low-affinity antibodies. Cell 186, 5486–5499 e5413. 10.1016/j.cell.2023.10.022.

28. Muramatsu, M., Kinoshita, K., Fagarasan, S., Yamada, S., Shinkai, Y., and Honjo, T. (2000). Class switch recombination and hypermutation require activation-induced cytidine deaminase (AID), a potential RNA editing enzyme. Cell 102, 553–563. 10.1016/s0092-8674(00)00078-7.

29. McKean, D., Huppi, K., Bell, M., Staudt, L., Gerhard, W., and Weigert, M. (1984). Generation of antibody diversity in the immune response of BALB/c mice to influenza virus hemagglutinin. Proc Natl Acad Sci U S A 81, 3180–3184. 10.1073/pnas.81.10.3180.

30. Jacob, J., Przylepa, J., Miller, C., and Kelsoe, G. (1993). In situ studies of the primary immune response to (4-hydroxy-3-nitrophenyl)acetyl. III. The kinetics of V region mutation and selection in germinal center B cells. J Exp Med 178, 1293–1307. 10.1084/jem.178.4.1293.

31. Jacob, J., and Kelsoe, G. (1992). In situ studies of the primary immune response to (4-hydroxy-3-nitrophenyl)acetyl. II. A common clonal origin for periarteriolar lymphoid sheath-associated foci and germinal centers. J Exp Med 176, 679–687. 10.1084/jem.176.3.679.

32. Di Noia, J.M., and Neuberger, M.S. (2007). Molecular mechanisms of antibody somatic hypermutation. Annu Rev Biochem 76, 1–22. 10.1146/annurev.biochem.76.061705.090740.

33. Goossens, T., Klein, U., and Kuppers, R. (1998). Frequent occurrence of deletions and duplications during somatic hypermutation: implications for oncogene translocations and heavy chain disease. Proc Natl Acad Sci U S A 95, 2463–2468. 10.1073/pnas.95.5.2463.

34. Mayer, C.T., Gazumyan, A., Kara, E.E., Gitlin, A.D., Golijanin, J., Viant, C., Pai, J., Oliveira, T.Y., Wang, Q., Escolano, A., et al. (2017). The microanatomic segregation of selection by apoptosis in the germinal center. Science 358. 10.1126/science.aao2602.

35. Stewart, I., Radtke, D., Phillips, B., McGowan, S.J., and Bannard, O. (2018). Germinal Center B Cells Replace Their Antigen Receptors in Dark Zones and Fail Light Zone Entry when Immunoglobulin Gene Mutations are Damaging. Immunity 49, 477–489 e477. 10.1016/j.immuni.2018.08.025.

36. Liu, Y.J., Joshua, D.E., Williams, G.T., Smith, C.A., Gordon, J., and MacLennan, I.C. (1989). Mechanism of antigen-driven selection in germinal centres. Nature 342, 929–931. 10.1038/342929a0.

37. Anderson, S.M., Khalil, A., Uduman, M., Hershberg, U., Louzoun, Y., Haberman, A.M., Kleinstein, S.H., and Shlomchik, M.J. (2009). Taking advantage: high-affinity B cells in the germinal center have lower death rates, but similar rates of division, compared to low-affinity cells. J Immunol 183, 7314–7325. 10.4049/jimmunol.0902452.

38. Shlomchik, M.J., Luo, W., and Weisel, F. (2019). Linking signaling and selection in the germinal center. Immunol Rev 288, 49–63. 10.1111/imr.12744.

39. Takeuchi, O., Fisher, J., Suh, H., Harada, H., Malynn, B.A., and Korsmeyer, S.J. (2005). Essential role of BAX,BAK in B cell homeostasis and prevention of autoimmune disease. Proc Natl Acad Sci U S A 102, 11272–11277. 10.1073/pnas.0504783102.

40. Nojima, T., Reynolds, A.E., Kitamura, D., Kelsoe, G., and Kuraoka, M. (2020). Tracing Self-Reactive B Cells in Normal Mice. J Immunol 205, 90–101. 10.4049/jimmunol.1901015.

41. Youle, R.J., and Strasser, A. (2008). The BCL-2 protein family: opposing activities that mediate cell death. Nat Rev Mol Cell Biol 9, 47–59. 10.1038/nrm2308.

42. Czabotar, P.E., Lessene, G., Strasser, A., and Adams, J.M. (2014). Control of apoptosis by the BCL-2 protein family: implications for physiology and therapy. Nat Rev Mol Cell Biol 15, 49–63. 10.1038/nrm3722.

43. Wei, M.C., Zong, W.X., Cheng, E.H., Lindsten, T., Panoutsakopoulou, V., Ross, A.J., Roth, K.A., MacGregor, G.R., Thompson, C.B., and Korsmeyer, S.J. (2001). Proapoptotic BAX and BAK: a requisite gateway to mitochondrial dysfunction and death. Science 292, 727–730. 10.1126/science.1059108.

44. Kuraoka, M., Yeh, C.H., Bajic, G., Kotaki, R., Song, S., Windsor, I., Harrison, S.C., and Kelsoe, G. (2022). Recall of B cell memory depends on relative locations of prime and boost immunization. Sci Immunol 7, eabn5311. 10.1126/sciimmunol.abn5311.

45. Motamedi, M., Xu, L., and Elahi, S. (2016). Correlation of transferrin receptor (CD71) with Ki67 expression on stimulated human and mouse T cells: The kinetics of expression of T cell activation markers. J Immunol Methods 437, 43–52. 10.1016/j.jim.2016.08.002.

46. Kepler, T.B. (2013). Reconstructing a B-cell clonal lineage. I. Statistical inference of unobserved ancestors. F1000Res 2, 103. 10.12688/f1000research.2-103.v1.

47. Smith, K.G., Weiss, U., Rajewsky, K., Nossal, G.J., and Tarlinton, D.M. (1994). Bcl-2 increases memory B cell recruitment but does not perturb selection in germinal centers. Immunity 1, 803–813. 10.1016/s1074-7613(94)80022-7.

48. Takahashi, Y., Cerasoli, D.M., Dal Porto, J.M., Shimoda, M., Freund, R., Fang, W., Telander, D.G., Malvey, E.N., Mueller, D.L., Behrens, T.W., and Kelsoe, G. (1999). Relaxed negative selection in germinal centers and impaired affinity maturation in bcl-xL transgenic mice. J Exp Med 190, 399–410. 10.1084/jem.190.3.399.

49. Boulianne, B., Rojas, O.L., Haddad, D., Zaheen, A., Kapelnikov, A., Nguyen, T., Li, C., Hakem, R., Gommerman, J.L., and Martin, A. (2013). AID and caspase 8 shape the germinal center response through apoptosis. J Immunol 191, 5840–5847. 10.4049/jimmunol.1301776.

50. Sugimoto-Ishige, A., Harada, M., Tanaka, M., Terooatea, T., Adachi, Y., Takahashi, Y., Tanaka, T., Burrows, P.D., Hikida, M., and Takemori, T. (2021). Bim establishes the B-cell repertoire from early to late in the immune response. Int Immunol 33, 79–90. 10.1093/intimm/dxaa060.

51. Meyer-Hermann, M.E., Maini, P.K., and Iber, D. (2006). An analysis of B cell selection mechanisms in germinal centers. Math Med Biol 23, 255–277. 10.1093/imammb/dql012.

52. Nakagawa, R., Toboso-Navasa, A., Schips, M., Young, G., Bhaw-Rosun, L., Llorian-Sopena, M., Chakravarty, P., Sesay, A.K., Kassiotis, G., Meyer-Hermann, M., and Calado, D.P. (2021). Permissive selection followed by affinity-based proliferation of GC light zone B cells dictates cell fate and ensures clonal breadth. Proc Natl Acad Sci U S A 118. 10.1073/pnas.2016425118.

53. Salerno, F., Whale, A.J., Matheson, L.S., Vespasiani, D., Foster, W.S., Mitchell, T.J., Screen, M., Stammers, M., Bell, S.E., Hodson, D.J., et al. (2025). RNA binding proteins control the G(2)-M checkpoint of the germinal center B cell. Sci Immunol 10, eadu3718. 10.1126/sciimmunol.adu3718.

54. Phan, R.T., and Dalla-Favera, R. (2004). The BCL6 proto-oncogene suppresses p53 expression in germinal-centre B cells. Nature 432, 635–639. 10.1038/nature03147.

55. Liu, M., Duke, J.L., Richter, D.J., Vinuesa, C.G., Goodnow, C.C., Kleinstein, S.H., and Schatz, D.G. (2008). Two levels of protection for the B cell genome during somatic hypermutation. Nature 451, 841–U811. 10.1038/nature06547.

56. Phan, R.T., Saito, M., Basso, K., Niu, H., and Dalla-Favera, R. (2005). BCL6 interacts with the transcription factor Miz-1 to suppress the cyclin-dependent kinase inhibitor p21 and cell cycle arrest in germinal center B cells. Nat Immunol 6, 1054–1060. 10.1038/ni1245.

57. Young, C., and Brink, R. (2021). The unique biology of germinal center B cells. Immunity 54, 1652–1664. 10.1016/j.immuni.2021.07.015.

58. Long, Z., Phillips, B., Radtke, D., Meyer-Hermann, M., and Bannard, O. (2022). Competition for refueling rather than cyclic reentry initiation evident in germinal centers. Sci Immunol 7, eabm0775. 10.1126/sciimmunol.abm0775.

59. Butt, D., Chan, T.D., Bourne, K., Hermes, J.R., Nguyen, A., Statham, A., O’Reilly, L.A., Strasser, A., Price, S., Schofield, P., et al. (2015). FAS Inactivation Releases Unconventional Germinal Center B Cells that Escape Antigen Control and Drive IgE and Autoantibody Production. Immunity 42, 890–902. 10.1016/j.immuni.2015.04.010.

60. Zhang, J., Kodali, S., Chen, M., and Wang, J. (2020). Maintenance of Germinal Center B Cells by Caspase-9 through Promotion of Apoptosis and Inhibition of Necroptosis. J Immunol 205, 113–120. 10.4049/jimmunol.2000359.

61. Pae, J., Schwan, N., Ottino-Loffler, B., DeWitt, W.S., Garg, A., Bortolatto, J., Vora, A.A., Shen, J.J., Hobbs, A., Castro, T.B.R., et al. (2025). Transient silencing of hypermutation preserves B cell affinity during clonal bursting. Nature 641, 486–494. 10.1038/s41586-025-08687-8.

62. Merkenschlager, J., Pyo, A.G.T., Silva Santos, G.S., Schaefer-Babajew, D., Cipolla, M., Hartweger, H., Gitlin, A.D., Wingreen, N.S., and Nussenzweig, M.C. (2025). Regulated somatic hypermutation enhances antibody affinity maturation. Nature 641, 495–502. 10.1038/s41586-025-08728-2.

63. Yeh, C.H., Finney, J., Okada, T., Kurosaki, T., and Kelsoe, G. (2022). Primary germinal center-resident T follicular helper cells are a physiologically distinct subset of CXCR5(hi)PD-1(hi) T follicular helper cells. Immunity 55, 272–289 e277. 10.1016/j.immuni.2021.12.015.

64. Song, S., Li, H., Yeh, C.H., Watanabe, A., Kuraoka, M., Gao, H., Shen, X., LaBranche, C.C., Williams, W.B., Saunders, K.O., et al. (2026). Functional Convergence of Genetically Diverse B-Cell Receptors in Simian-HIV Infected Rhesus Macaques. bioRxiv. 10.64898/2026.01.09.698730.

65. Ioerger, T.R., Du, C., and Linthicum, D.S. (1999). Conservation of cys-cys trp structural triads and their geometry in the protein domains of immunoglobulin superfamily members. Mol Immunol 36, 373–386. 10.1016/s0161-5890(99)00032-2.

66. Tiller, T., Busse, C.E., and Wardemann, H. (2009). Cloning and expression of murine Ig genes from single B cells. J Immunol Methods 350, 183–193. 10.1016/j.jim.2009.08.009.

67. Dunbar, J., and Deane, C.M. (2016). ANARCI: antigen receptor numbering and receptor classification. Bioinformatics 32, 298–300. 10.1093/bioinformatics/btv552.

68. Delgado, J., Radusky, L.G., Cianferoni, D., and Serrano, L. (2019). FoldX 5.0: working with RNA, small molecules and a new graphical interface. Bioinformatics 35, 4168–4169. 10.1093/bioinformatics/btz184.

69. Ruffolo, J.A., Chu, L.S., Mahajan, S.P., and Gray, J.J. (2023). Fast, accurate antibody structure prediction from deep learning on massive set of natural antibodies. Nat Commun 14, 2389. 10.1038/s41467-023-38063-x.

